# NUCLEAR RETENTION OF LEPTIN/UPD2 REGULATES ORGANISMAL RESILIENCE TO NUTRIENT EXTREMES

**DOI:** 10.1101/2021.01.29.428913

**Authors:** Michelle E. Poling, Camille E. Sullivan, Aditi Madan, Kevin P. Kelly, Ava E. Brent, Julien Dubrulle, Prashant Raghavan, Akhila Rajan

## Abstract

Adipokines released from the adipocytes function as a systemic adipometer; they impinge on neural circuits to signal nutrient status. On starvation, adipokines must be retained to signal energy deficit; else, it significantly reduces starvation survival. But how fat cells retain adipokines is unclear. Here, we demonstrate that Atg8, a cell-intrinsic autophagy factor, regulates the starvation-induced acute retention of the Leptin *Drosophila* ortholog Upd2. We show that on starvation, as a direct consequence of Atg8’s lipidation, Upd2 accumulates in the nucleus. We illustrate that Upd2’s nuclear retention is critical to fat mobilization and increased starvation resilience. Furthermore, nuclear Upd2 promotes the expression of a secreted innate immune gene signature. This hints at an unanticipated connection between adipokine nuclear retention and increased innate immunity. In conclusion, we propose that, during starvation, Atg8’s role is not just limited to autophagy but is critical for withholding adipokines in the nucleus to promote starvation resilience.

**GRAPHICAL ABSTRACT:** 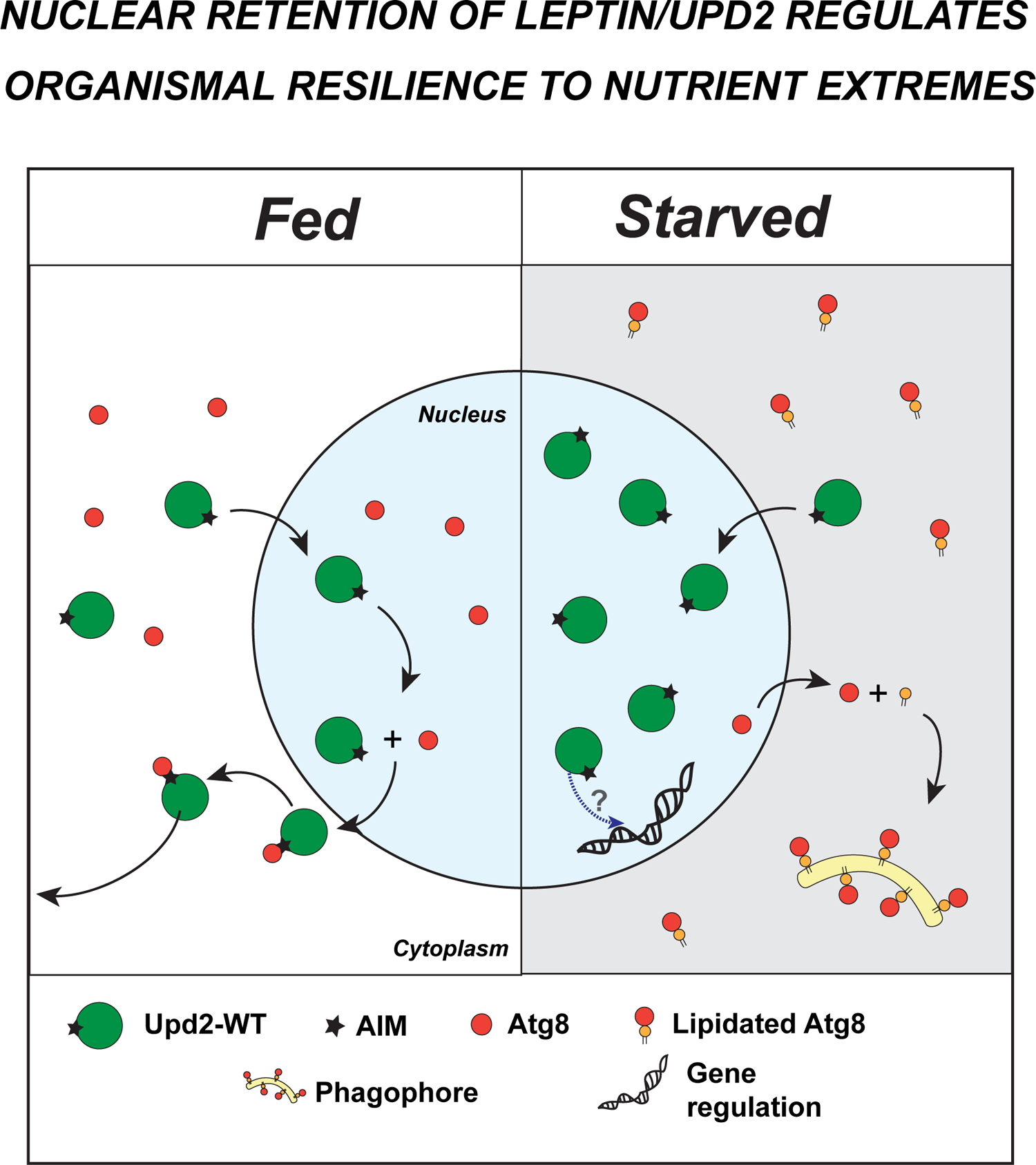

1. In fed state Upd2 requires Atg8 for nuclear exit and cytosolic localization.
2. Atg8’s lipidation on starvation results in Upd2’s nuclear accumulation.
3. Upd2 nuclear retention on starvation increases fat mobilization and post-starvation hunger.
4. On starvation Upd2 nuclear retention increases expression of a secreted innate immune signature.

## INTRODUCTION

Faced with constant fluctuations in the food supply, organisms rely on accurate nutrient sensing to maintain a steady internal milieu [1]. Regulation of nutrient-sensing can be broadly divided into two categories. One includes cell-intrinsic players such as kinases like AMPK, TOR [2] and molecular players of the nutrient-scavenging autophagy pathway [3]. Other factors that orchestrate systemic cell-extrinsic responses include the three key metabolic hormones, Insulin, Glucagon, and Leptin [4]. For robust energy homeostasis, cell-intrinsic and extrinsic mechanisms must tightly coordinate their roles; but the precise mechanisms underlying such coordination remain to be discovered.

The evolutionary of both cell-intrinsic nutrient-sensing [5, 6] and inter-organ energy homeostasis [7–9] between flies and mammals are now well-appreciated. In particular, this conservation includes both lipid sensing and its dysregulation, including the development of insulin resistance and ‘diabetic’ phenotypes [9–16]. Hence, discoveries in the *Drosophila* model regarding how endocrine and cell-intrinsic signals interact will broadly be relevant. In this study, using this *Drosophila* system, we investigated how a cell-intrinsic starvation sensing pathway acutely regulates one endocrine factor produced by the adipose tissue.

Adipocytes release hormones to relay fat store status systemically. Leptin in mammals [17, 18] and its functional ortholog in fruit flies, Unpaired2 (Upd2) [19], are primary adipokines that are released in proportion to fat stores [20–22]. Both in flies and mammals, Leptin/Upd2 impinges on brain circuits that control energy expenditure, appetite, and overall metabolism [19, 23–25]. Adipokines convey a permissive signal in surplus nutrient states, indicating that energy resources can be devoted to costly activities like immunity, sleep, and reproduction. During periods of scarcity, their circulation is reduced, signaling an energy deficit, and enabling organisms to conserve energy.

Starvation-induced reduction in Leptin/Upd2 levels is crucial for an organism’s neuroendocrine response to reduced energy reserves. Specifically, Leptin injections during fasting dysregulate neuroendocrine physiology and decrease survival capacity in mice [17]. Based on this study [17], Flier and colleagues have proposed that “*the primary physiologic role of Leptin is to provide a signal of energy deficit to the CNS*” [4, 26]. Similarly, flies with reduced levels of Upd2 display increased starvation survival [19, 27], suggesting that Upd2’s retention is critical to enable starvation resilience. But how starvation induces adipokine retention remains to be fully characterized.

In our previous work, we identified that chronic nutrient deprivation (> 5 days) results in a significant reduction in steady-state Upd2 mRNA levels [19]. We also showed that prolonged starvation (> 3 days) dismantles the secretion machinery utilized by Upd2 [27]. However, these retention mechanisms operate over a temporal span of days and are not in play within an acute time window (4-8 hours). Therefore, we wondered what other acute mechanisms are utilized by fat cells to retain adipokines.

Autophagy is an acute cell-intrinsic response to nutrient deprivation[28–30]. In adipocytes, autophagy controls lipid metabolism by a process known as lipophagy [29, 31] Autophagy-related protein-8 (Atg8) is a central player in the autophagy pathway. Atg8 (LC3/GABARAP in mammals) is a ubiquitin-like protein family member [32, 33]. Atg8 is widely distributed in the nucleus in fed cells [34]. It translocates to the cytoplasm during starvation, where it is conjugated to the lipid moiety phosphatidylethanolamine (PE)[35]. Lipidated Atg8 regulates autophagosome fusion to lysosomes, resulting in cargo degradation in the cytosol [36]. In addition to its well-studied role in autophagy, cytosolic Atg8 regulates other non-degradative cellular activities [37], including but not limited to phagocytosis [38], viral replication [39], extracellular vesicle secretion[40], and endocytosis[41].

In this study, we reveal a novel unconventional role for the nuclear pool of Atg8 in regulating nutrient-state dependent adipokine localization. In fed cells, Upd2 requires nuclear Atg8 for cytosolic localization. On starvation, Atg8 translocates to the cytosol and is lipidated; this depletes Atg8’s nuclear availability. As a direct consequence of Atg8’s lipidation, we show that Upd2 is withheld in the nucleus. We illustrate that Upd2’s nuclear accumulation promotes starvation resilience but increases sensitivity to high-sugar diets. Furthermore, we find that Upd2 nuclear retention upregulates a secreted innate immune gene signature. This hints at adipokine retention, promoting an anticipatory increase in innate immunity during periods of non-feeding. In sum, we uncover an unexpected link between Atg8 and adipokine signaling and illustrate how their interplay controls response to nutrient extremes.

## RESULTS

### ACUTE STARVATION TRIGGERS UPD2 NUCLEAR ACCUMULATION

To observe endogenous Upd2 localization in *Drosophila* adult adipocytes, we generated CRISPR-engineered tagged knock-ins into endogenous Upd2 genomic locus (See methods). The Upd2-GFP and Upd2-HA genomic knock-ins were able to rescue *upd2*-deletion (*upd2Δ*) fly fat storage defects (Figure S1A), indicating that endogenous genomic tagging (either with HA and GFP) preserved Upd2 function. In place of a robust antibody for immunostaining the Upd2 protein, HA or GFP staining of Upd2 endogenous tagged knock-ins allowed us to follow Upd2’s endogenous localization. We observed that within 4 hours of starvation, Upd2-GFP levels increased overall, but especially in the nucleus (Figure 1Ab, and 1A’). This increase in the Upd2 nuclear signal was also observed in the Upd2-HA tag genomic knock-in (Figure S1B), providing a second line of evidence that endogenous Upd2 levels in the adult fly fat nuclei are higher on starvation.

Using another cell system, we sought to corroborate the increased starvation-induced increase in Upd2’s nuclear localization that we observed in adult fly fat cells. *Drosophila* S2R+ cells are derived from embryonic stages and have macrophage-like properties [42]. *Drosophila* S2R+ cells respond robustly to amino-acid (AA) deprivation by inducing nutrient deprivation signals [43, 44]. Given this, we transiently transfected cells with Upd2-WT::GFP and assayed Upd2-WT::GFP localization in cells cultured in complete media (fed) of in amino-acid (AA) depleted media (starved). Akin to the increase in Upd2 nuclear accumulation in fly fat on acute starvation, we observed a significant increase in Upd2 nuclear accumulation (Figure 1Bb) within 4-8 hours of AA deprivation. Then, we asked whether Upd2’s nuclear accumulation was reversible within a short-time frame of refeeding. We noted that when AA-acid starved cells were cultured in complete media within 4-6 hours of culturing cells, Upd2::GFP localizes in punctate cytosolic structures (Figure 1Bc, Bd; See 1B’s for quantification). Collectively, studies from both S2R+ cells and fly fat suggested a strong direct correlation between starvation and Upd2 nuclear accumulation.

We wanted to determine if Upd2’s starvation-induced nuclear accumulation correlated with its release into extracellular space. Detection of endogenous Upd2 in the adult fly hemolymph, either by western or quantitative ELISA, is a long-standing technical hurdle. We and others have utilized the S2R+cell culture system to quantify Upd2’s release into the media reliably [27, 45, 46]. Moreover, genome-wide functional screens for protein secretion in S2R+ cells have identified genes relevant to secretion biology *in vivo* [47, 48]. Hence, given that findings from the *Drosophila* S2R+ system regarding secretion in general, and Upd2 secretion in particular, are recapitulated *in vivo*, we utilized this system to investigate the relationship between Upd2 nuclear accumulation and its secretory state.

We measured Upd2’s secretion in media of S2R+ cells cultured in complete versus AA-depleted media using a quantitative ELISA for Upd2::GFP (See methods). AA deprivation, within 6 hours, reduced Upd2 secretion by 76%. Notably, Upd2 was restored to normal levels within 5 hours (Figure 1C) when cells were cultured in complete media. We noted the negative correlation between Upd2’s nuclear accumulation (4-hours of starvation) and its secretory status. This led us to hypothesize that acute nuclear retention impedes Upd2 secretion. Accordingly, we observed that the knockdown of *Embargoed* (*Emb*) [49], the *Drosophila* CRM1 nuclear export factor, causes increased Upd2 nuclear retention (Figure S1C) and impedes Upd2 secretion (Figure S1D). We cannot exclude the possibility that a blunt disruption to nuclear export will likely impact secretion globally. Nonetheless, we noted that Upd2’s nuclear accumulation inversely correlated with its secretory potential. Altogether, we concluded that acute starvation reduces Upd2’s extracellular release and increases Upd2 nuclear accumulation in diverse contexts of *Drosophila* S2R+ cells and adult fly fat.

**Figure 1.**
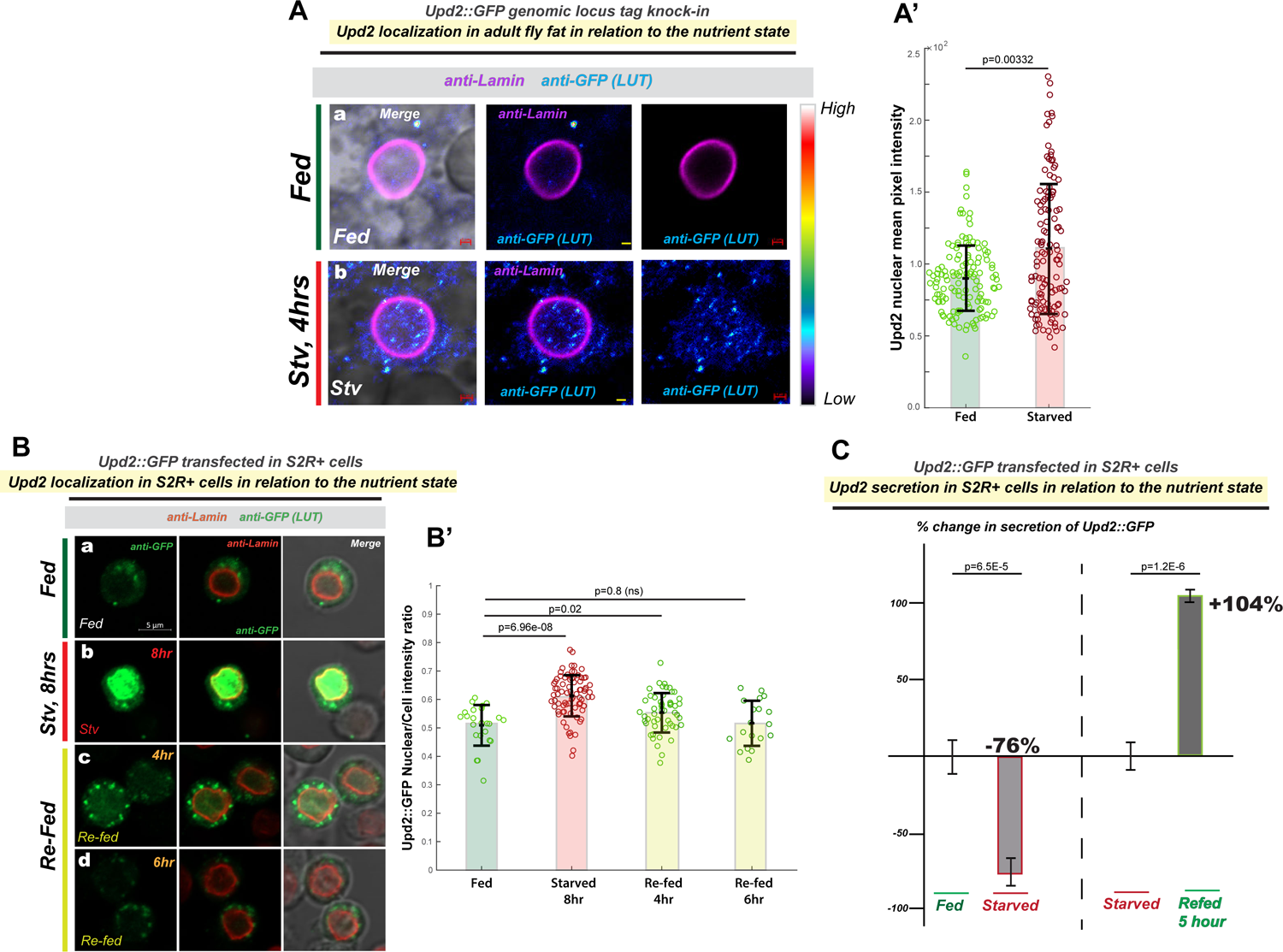
Acute Starvation triggers Upd2 nuclear accumulation. (A) <u>Effect of starvation on Upd2 nuclear accumulation in adult fly fat tissue:</u> Confocal micrographs of fixed and immunostained adult fly fat from flies CRISPR-engineered (see methods) with GFP tag in the endogenous Upd2 locus (that rescues the upd2Δ mutant- see Figure S1). Fat is stained with GFP antibody (in a, b) in well-fed flies (a), or flies starved for 4 hours. Upd2 shows increased nuclear accumulation on starvation [see look up table (LUT)]. Scale bar is 1µM. (A’) Upd2-GFP endogenous nuclear accumulation is quantified using 3D-volumetric segmentation-based calculations (See Methods). Each dot represents a fat cell nucleus, 75-100 fat nuclei were counted per condition per genotype. Statistical significance is quantified by the two-sided Wilcoxon test. Also see Figure S1 for similar experiment performed on an HA-tag knock into the Upd2 genomic locus (Upd2-HA). (B) <u>Effect of starvation and re-feeding on Upd2’s nuclear accumulation in *Drosophila* S2R+ cells:</u> Confocal micrographs of single optical-slices of *Drosophila* S2R+ cells transiently transfected with Upd2-WT::GFP (green; anti-GFP) and co-stained with Lamin (red). Scale bar is 5um. In B’, the ratio of Upd2::GFP nuclear/whole cell intensity is plotted. Each dot represents a cell, 50-100 cells were counted per time point. Statistical significance is quantified by the two-sided Wilcoxon test. (C) <u>Effect of starvation and re-feeding on Upd2’s release into media:</u> Normalized percent fold change in secreted GFP signal detected by GFP sandwich ELISA assay performed on conditioned media of S2R+ cells transiently transfected with Upd2-WT::GFP. Cells were incubated either with complete media (control) or Amino-Acid (AA) free media for the indicated times. The refeeding % fold change is indicated with respect to starvation. Statistical significance is quantified by unpaired two-tailed t-test on 6 biological replicates per condition.

### UPD2 REQUIRE S ATG8 FOR NUCLEAR EXIT

Intriguingly, although Upd2 required CRM1/Emb-based nuclear export (Figure S1D), we were unable to identify a canonical nuclear export signal (NES) on Upd2, suggesting that Upd2’s nuclear export occurs indirectly via other protein adapters (See Discussion). To identify clues to what Upd2’s protein partners are likely to be, we examined protein motifs in Upd2’s protein sequence. We noted that Upd2 had several Atg8-interaction motifs (AIM) [50]. Given Atg8’s well-characterized role in nutrient-sensing, we wondered whether Atg8 might interact with Upd2 to control its nutrient-dependent nucleocytoplasmic localization.

We generated transgenic flies with two-point mutations to Upd2’s putative AIM-like sequence (Figure 2A). Multiple AIM-like sequences are found in Upd2, but only one is the canonical ‘WXXL’ AIM[50]; hence we mutated that. We specifically expressed Upd2-AIM-(WXXL→AXXA) in fly fat tissue in an *upd2*-deletion background (*upd2Δ; Lpp-Gal4> UAS-Upd2-AIM-::mCherry*) and examined Upd2’s localization in the abdominal fat pads of 7 day-old adult male flies, fed *ad libitum* in relation to control transgene (*upd2Δ; Lpp-Gal4> UAS-Upd2-WT::mCherry*). Upd2-AIM-expressing flies displayed a higher nuclear accumulation (2Ab) of Upd2 than controls (2Aa). It is to be noted that the Upd2-WT and Upd2-AIM-transgenes are expressed at similar levels (Figure S7B). Furthermore, corroborating the role of Atg8 in regulating Upd2’s localization, fat tissue-specific, acute temporal knockdown of Atg8 (*upd2GFP-knockin; ppl-Gal4, tubGal80^ts^> Atg8-RNAi*) resulted in a significant increase in the endogenous Upd2’s mean fly fat nuclear intensity (Figure 2B). Thus, disruption of Upd2’s putative AIM or knockdown of Atg8 increases Upd2’s nuclear accumulation and phenocopies acute starvation.

**Figure 2.**
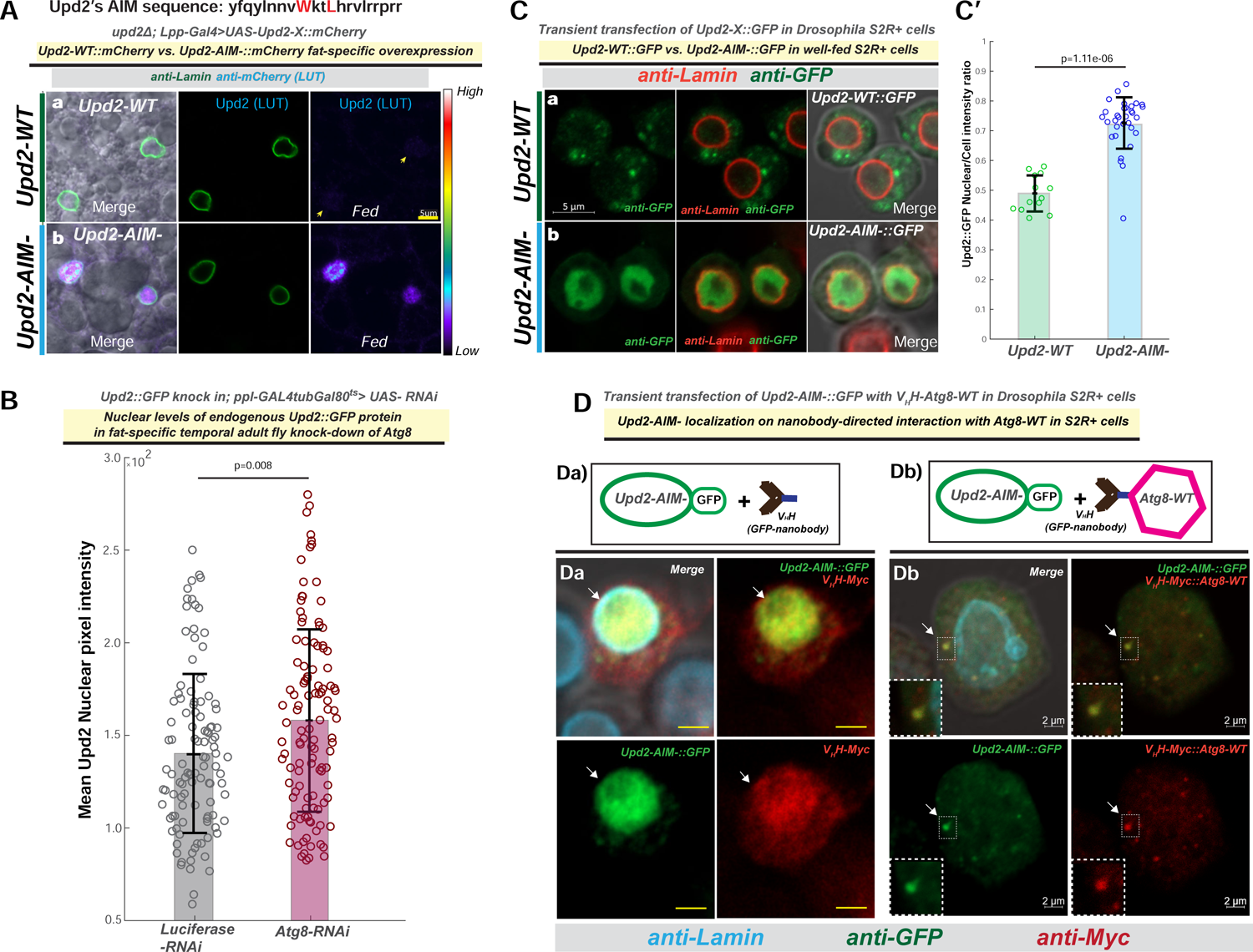
Atg8 controls Upd2 nucleocytoplasmic localization. (A) <u>Effect of point mutations to Upd2’s AIM (Upd2-AIM-) on its nuclear accumulation in adult fly fat:</u> Upd2-AIM-includes two point mutations to Upd2’s putative AIM (WXXL→ AXXA) – see sequence above the figure. Confocal micrographs of single optical sections of fixed and immunostained adult fly fat from transgenic flies expressing mCherry tagged Upd2-cDNAs for (a) Upd2-WT (UAS-Upd2-WT) or (b) Upd2’s AIM mutated (UAS-Upd2-AIM-) under the control of a fat-cell specific promoter (Lpp-Gal4) in a upd2-deletion background (upd2Δ; Lpp-Gal4). (B) <u>Effect of Atg8-KD on Upd2 nuclear accumulation in adult fly fat tissue:</u> Endogenous Upd2 nuclear signal in fly fat (upd2::GFP-endogenous knock-in) of flies expressing control (Luciferase-grey bar) or Atg8-RNAi (red bar) in fat cells during adult stages (ppl-Gal4tubGal80ts). (C) <u>Effect of point mutations to Upd2’s AIM (Upd2-AIM-::GFP) on its nuclear accumulation in *Drosophila*</u> <u>S2R+ cells:</u> Confocal micrographs (a, b) of single optical-slices of *Drosophila* S2R+ cells transiently co-transfected with (a) Upd2-WT::GFP or ( b) Upd2-AIM-::GFP (green-anti-GFP) stained with Lamin (red) in a fed state. Scale bars are 5um for a, b. In C’, the ratio of Upd2::GFP nuclear/whole cell intensity is plotted. See methods of details on quantification. For (B) each dot on the graph is a fat cell nucleus, in (C’) each dot is a cell. Statistical significance for B and C’ are quantified by the two-sided Wilcoxon Rank sum tests. (D) <u>Effect of nanobody-directed interaction between Atg8-WT and Upd2-AIM-::GFP on Upd2’s localization</u> <u>in *Drosophila* S2R+ cells:</u> Top panel shows schematic of the experimental design. V_H_H tag, a small 15kDa antibody to GFP, is expressed by itself (a) or fused to the N-terminal of Atg8-WT and co-transfected with Upd2-AIM-::GFP. Bottom panel shows a single XY-slice of confocal image in *Drosophila* S2R+ cells cultured in complete media. Cells were fixed and immunostained with antibodies to detect V_H_H-Myc / V_H_H-Myc::Atg8-WT (anti-Myc-red) and Upd2-AIM-::GFP (anti-GFP). In a, note that Upd2-AIM-::GFP displays nuclear accumulation and so does V_H_H-Myc, consistent with its role as an antibody to GFP (see arrows). In b, punctate overlap between V_H_H Atg8 and GFP are observed-see arrow and inset. Note Upd2-AIM- is present in the cytosolic puncta, instead of nuclear accumulation (compare with Figure 2Cb). Also see Figure 3B’ for quantification of the nuclear/cytosolic localization of Upd2-AIM- in the presence of V_H_H-Myc::Atg8-WT compared to control (V_H_H-Myc empty).

Next, we wanted to test whether Upd2’s intracellular localization in *Drosophila* S2R+ cells, like what we observe for fly fat, is regulated by Atg8. In Drosophila S2R+ cells, we observed that Upd2-AIM-expression was significantly nuclear (Figure 2Cb, C’) and recapitulated the Upd2-AIM-localization in fly fat (Figure 2Ab). Co-immunoprecipitation (Co-IP) experiments, performed in *Drosophila* S2R+ cells, revealed that Upd2-WT could be complex with Atg8, but point-mutations to Upd2’s putative AIM (WXXL→ AXXA) impair its ability to complex with Atg8 (Figure S2A). Though these experiments are performed in supraphysiological concentrations, the co-IP complex observations indicate that Upd2 and Atg8 are likely to be complex and that Upd2’s AIM sequence is required for this interaction. Furthermore, corroborating Atg8’s requirement for Upd2’s cytosolic localization, Atg8 knockdown in *Drosophila* S2R+ cells, using two independent dsRNAs, resulted in a significant increase in Upd2-WT::GFP nuclear signal (Figure S2B). These observations lead us the hypothesize that Atg8 regulates Upd2’s cytosolic localization during a fed state.

Atg8 knockdown (Figure S2B), or point-mutations to Upd2’s AIM (Figure 2C), cause increased Upd2 nuclear accumulation in S2R+ cells. Prior observations suggested that Upd2 nuclear accumulation inversely correlates with its secretory state (Figure 1C, S1E). When we performed quantitative ELISA assays for Upd2 secretion during an Atg8-knockdown (KD) or in Upd2-AIM-state, we observed a significant impairment of Upd2 secretion (Figure S2C and S2D). A previous study identified that Upd2 undergoes GRASP-mediated unconventional secretion[27]. In keeping with a role for Atg8 in mediating unconventional GRASP-mediated secretion [40, 51–53], we found using Co-IP experiments in S2R+ cells that Upd2-AIM-ability to interact with GRASP is impaired. In sum, these experiments suggested that Upd2, via its AIM sequence, interacts with Atg8, and this interaction regulates both its cytosolic localization and subsequent extracellular release in fed cells.

Finally, we sought to test whether reconstituting Upd2-AIM-interaction with Atg8 would be sufficient to localize Upd2-AIM- to the cytosol. To test this, we took advantage of the genetically encoded protein binder tag V_H_H that acts as a nanobody recognizing GFP (vhhGFP4) [54]. Functional studies performed by fusing V_H_H to different proteins and then asking how it affects the localization of a GFP-tagged protein have been used to dissect molecular players controlling protein localization [55–57]. Akin to the experimental design in the studies mentioned above, which utilized nanobody fusions to probe protein localization, we tested whether the interaction between Upd2-AIM-::GFP and VHH tagged was reconstituted Atg8 is sufficient to restore the cytosolic localization of Upd2-AIM- in fed cells (See Schematic in Figure 2D).

To this end, in *Drosophila* S2R+ cells, we co-expressed with Upd2-AIM-::GFP either the control V_H_H::Myc (control) or Atg8-WT was fused to V_H_H::Myc (V_H_H::Myc-Atg8-WT). As would be expected of a GFP-nanobody, in that it would bind to GFP-tagged proteins, we observed that V_H_H::Myc showed increased nuclear accumulation, overlapping with Upd2-AIM-::GFP (See arrow; Figure 2Da). However, co-expression of the control nanobody did not result in localization change of Upd2-AIM-::GFP. Strikingly, in the presence of V_H_H-tagged Atg8-WT (V_H_H::Myc-Atg8-WT), we observed that Upd2-AIM-GFP is now cytosolic and co-localized with Atg8-WT in punctate structures (See Arrow; Figure 2Db), and reflecting this, Upd2-AIM-nuclear accumulation was significantly reduced (See quantification in Figure 3B’). Hence, reconstituting the physical interaction between Atg8 and Upd2-AIM- is sufficient to localize Upd2-AIM- to the cytosol. This pivotal result demonstrated that Upd2 requires Atg8 for its nuclear exit and cytosolic localization.

**Figure 3.**
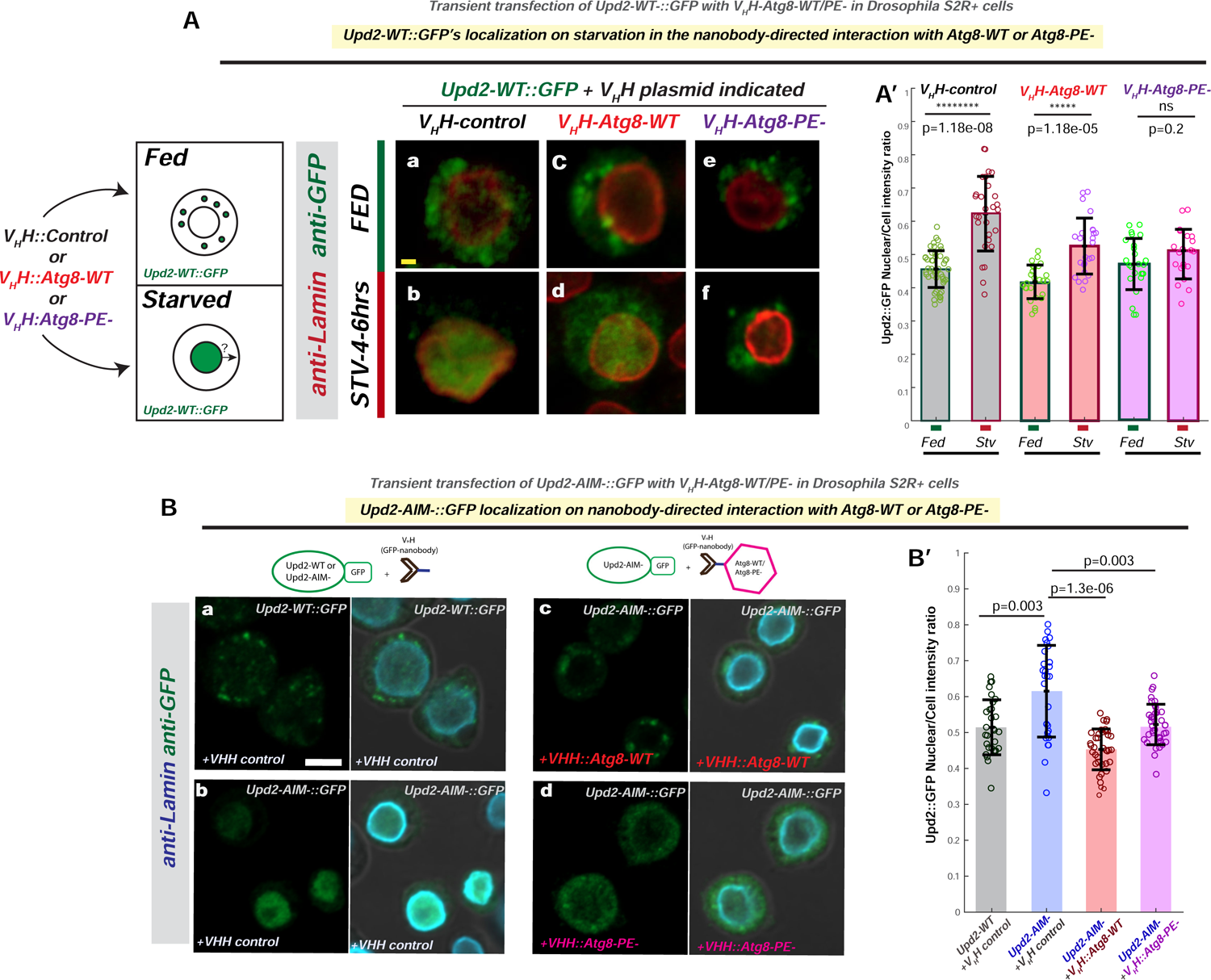
Atg8 lipidation status regulates starvation-induced nuclear retention. In (A) and (B), schematics illustrate experimental design. V_H_H tag, a small 15kDa antibody to GFP, is the control and V_H_H tag fused to the N-terminal of Atg8-WT or Atg8-G◊A mutation (Atg8-PE-) Also see companion Figure S3 and S4. (A) <u>Effect of nanobody-directed interaction between Upd2-WT::GFP and Atg8-WT or Atg8-PE-in</u> *<u>Drosophila</u>* <u>S2R+ cells during an acute amino-acid starvation:</u> Side panels show schematic of the experimental design. Upd2’s starvation induced nuclear retention is assessed. Aa-Af show a single XY-slice of confocal image in *Drosophila* S2R+ cells cultured in complete media (a, c, e) or amino-acid free media (b, d, f). Cells were fixed and immunostained with anti-Lamin (blue) and anti-GFP (green) to detect Upd2-WT::GFP. Scale bar represents 1µM. (B) <u>Effect of nanobody-directed interaction between Upd2-AIM-::GFP and Atg8-WT or lipidation defective</u> <u>Atg8-PE-in *Drosophila* S2R+ cells:</u> Top panels show schematic of the experimental design. Ba-Bd show a single XY-slice of confocal image in *Drosophila* S2R+ cells cultured in complete media. Cells were fixed and immunostained with anti-Lamin (blue) and anti-GFP (green) to detect Upd2-WT::GFP (a) or Upd2-AIM-::GFP (b-c). The effect of co-transfection with of Upd2-WT with V_H_H tag (a) is used as baseline and compared to Upd2-AIM-localization in the presence of ‘empty’ V_H_H tag (b) or V_H_H tag fused to Atg8-WT (c) or Atg8-PE-(d) is assessed. Scale bar represent 5µM. A’ and B’ graphs show 3D-volumetric quantification of nuclear to whole cell intensity of GFP signal. Statistical significance is calculated using two-sided Wilcoxon sum rank tests. Each circle represents a cell.

### ATG8’S LIPIDATION CAUSES STARVATION-INDUCED UPD2 NUCLEAR RETENTION

We were puzzled, however, that despite the presence of endogenous Atg8, on starvation Upd2 nuclear retention phenocopies Atg8’s knockdown and Upd2-AIM-. We reasoned that, on acute starvation, Atg8’s recruitment to cytosolic vesicles might preclude Atg8 from participating in Upd2’s nuclear exit. In line with this hypothesis, pharmacological activation of autophagy, by treatment with autophagy activator Torin [58], results in a dose-dependent increase in Upd2 nuclear accumulation (Figure S3); this indicated that increased autophagy increases Upd2 nuclear retention. Torin treatment increases Atg8’s lipidation [59]. Since lipidated Atg8 associates with cytosolic membrane structures [36, 60], this prevents Atg8 from freely diffusing between the nucleus and cytosol [61].

We posited that Atg8’s lipidation is likely to reduce its nuclear availability; consequently, Upd2’s nuclear exit is impeded, resulting in Upd2’s nuclear accumulation. To test this hypothesis, we generated lipidation-defective Atg8 by mutating to the C-terminal lipid acceptor moiety glycine of Atg8 (Atg8-G→A) [35]. We performed tests on Atg8 G→A to validate that lipidation was defective by using accepted assays in the field to test Atg8 lipidation [62]. This included: i) doublet on western blots of starvation lysates (Figure S4A); ii) recruitment to cytosolic vesicular structures on starvation (Figure S4B). From this, we inferred that Atg8 G→A mutation rendered its lipidation defective (hereafter, it is referred to as Atg8-PE-for simplicity). We fused the lipidation defective Atg8-PE- to an N-terminal V_H_H tag ( V_H_H::Myc-Atg8-PE-; see methods). In subsequent experiments, we utilized V_H_H::Myc-Atg8-PE- to test the role of Atg8 lipidation in Upd2’s starvation-induced nuclear accumulation.

We hypothesized that lipidation of Atg8 reduces its capacity to participate in Upd2 nuclear exit. Hence, we predicted that reconstituting Upd2-WT::GFP’s interaction with lipidation-defective Atg8 should ‘rescue’ its nuclear accumulation defect (Figure 3A- see schematic). Consistent with this prediction, we found that, when co-transfected with lipidation defective Atg8-PE-(V_H_H::Myc-Atg8-PE-), Upd2 is cytosolic even on starvation (Figure 3Af, 3A’). Furthermore, in support of our hypothesis, lipidation competent Atg8-WT (V_H_H::Myc-Atg8-WT) does not reduce starvation-induced Upd2 nuclear accumulation (Figure 3Ad; A’). Studies have shown that Atg8/LC3 lipidation increased bulk preventing Atg8/LC3 from shuttling between the nucleus and cytosol [61]. Accordingly, we observe that Atg8-WT is localized to cytosolic structures on starvation (Figure S4Bb), but Atg8-PE-does not (Figure S4Bd). Collectively, these experiments (Figure 3A) suggest that, on starvation, Atg8’s lipidation reduces its nuclear availability, in turn resulting in increased Upd2 nuclear accumulation.

This observation also suggested that Atg8’s lipidation is not required for Upd2’s nuclear exit in fed cells. Accordingly, V_H_H nanobody-based reconstitution of Upd2-AIM-::GFP with Atg8-PE-(Figure 3Bd), similar to Atg8-WT (Figure 3Bc), is sufficient to re-localize Upd2-AIM-::GFP to the cytosol (Figure 3B, B’). Taken together, these results (Figure 3) strongly suggest that Atg8’s lipidation, on starvation, recruits it away from the nucleus to the cytosol. As a direct consequence, Upd2 is withheld in the nucleus.

### ATG8’S LIPIDATION IN *DROSOPHILA* ADULT FAT REGULATES OUTCOME ON STARVATION CONTINGENT ON UPD2

The nuclear to cytosolic translocation of Atg8 on starvation is a key step of regulation in the lipidation of Atg8 [34]. Therefore, we predicted that consistent with mammalian data, our results in *Drosophila* S22R+ cells (Figure S4B), Atg8’s nuclear amount *in vivo* in adult fly fat cells will be depleted on starvation. Flies expressing GFP tagged Atg8 in fly fat (*Lpp-Gal4> UAS-GFP::Atg8-WT*) were subjected to an acute starvation diet (4-6 hours 0% sucrose agar) or *ad libitum* fed 30% high sugar diet (HSD) for 14 days. We assessed the total nuclear signal of GFP::Atg8-WT in 3D volume between the flies fed *ad libitum* normal lab food versus starvation and HSD (Figure 4A; Also see Study Design section). As predicted, acute starvation significantly reduced the total nuclear signal of GFP::Atg8-WT, in conjunction with an increase in cytosolic punctate GFP::Atg8-WT (Figure 4Ac, 4A’). Conversely, a surplus diet (30% high-sugar) significantly increased the total nuclear signal of GFP::Atg8-WT (Figure 4Ab, 4A’). In sum, in *Drosophila* adult fly fat Atg8, nuclear signal increases on a surplus diet, whereas Atg8 is less nuclear on starvation.

**Figure 4.**
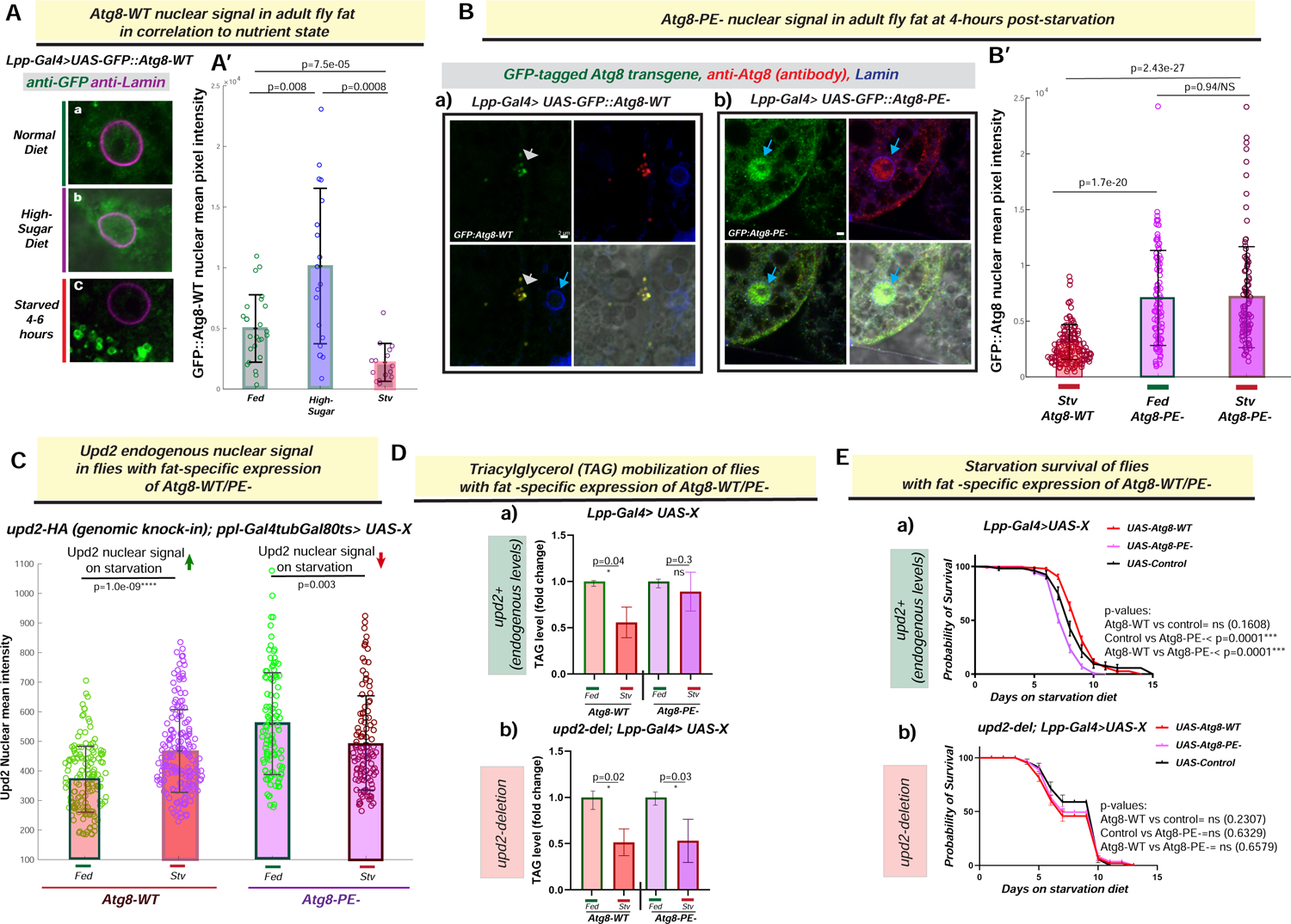
Effects of lipidation-defective Atg8 expression in fly fat in relation to Upd2. (A) <u>Atg8-WT::GFP nuclear levels in fly fat correlate with nutrient state:</u> Confocal micrographs of XY-slices from fixed and immunostained adult fly fat from flies expressing GFP::Atg8-WT transgene stained with anti-GFP (green) to detect Atg8. Atg8’s nuclear localization in three different nutrient states-normal lab food (a), 14-day high sugar diet (b) and acute starvation (c) are shown. (B) <u>Lipidation defective Atg8 (Atg8-PE-) remains nuclear in *Drosophila* adult fly fat on acute starvation:</u> Confocal micrographs of single optical sections of fixed and immunostained [anti-GFP(green); anti-Atg8 (red); anti-Lamin (blue)] adult fly fat from transgenic flies-starved for 4 hours-expressing GFP-tagged Atg8-cDNAs for (Da) Atg8-WT (*UAS-GFP::Atg8-WT*) or (b) lipidation-defective Atg8 (*UAS-GFP::Atg8-PE-*) under the control of a fat-cell specific promoter (*Lpp-Gal4*) in an Atg8-WT background. Scale bar represents 2µM. In A’ and B’ the amount of GFP::Atg8 in the nucleus in quantified in using 3D-volumteric image segmentation (see methods). Each circle represents a nucleus and statistical significance is calculated using two-sided Wilcoxon sum rank tests. (C) <u>Effect of fat-specific expression of Atg8-WT/PE- on Upd2 nuclear accumulation in adult fly fat tissue:</u> Upd2-GFP endogenous nuclear accumulation is quantified using 3D-volumetric segmentation-based calculations in flies with acute fat-specific expression of Atg8-WT or Atg8-PE- in fed versus 4 hours starvation. Each dot represents a fat cell nucleus, 75-100 fat nuclei were counted per condition per genotype. Statistical significance is quantified by the two-sided Wilcoxon test. For D and E, Flies expressing cDNAs (UAS-X) for GFP::Atg8-WT, GFP::Atg8-PE- and control (*UAS-Luciferase*), specifically in fat tissue (*Lpp-Gal4>UAS-X*), with endogenous levels of Upd2 (a) or upd2-deletion (*upd2Δ; Lpp-Gal4>UAS-X*). (D) <u>Effect of fat-specific expression of Atg8-PE- on triglyceride mobilization during starvation in the</u> <u>presence and absence of endogenous Upd2:</u> TAG levels at fed state were held at 1, and fold change in TAG at starvation was calculated. Error bars represent standard deviation of 9 biological replicates. Statistical significance is calculated using the two-tailed student’s T test. (E) <u>Effect of fat-specific expression of Atg8-PE- on starvation survival in the presence and absence of</u> <u>endogenous Upd2:</u> Probability of survival of transgenic flies with fat-specific overexpression of Atg8-PE-in a WT (a) and upd2 deletion background (b) are shown. Statistical significance is calculated using the Log-rank (Mantel-Cox) test. Per genotype >100 flies were used per experiment. Data shown is then consolidation of three independent experiments.

Next, we wondered what happens to nuclear levels of lipidation defective Atg8 in adult fly fat cells on starvation. Whereas the GFP::Atg8-WT (Figure 4Ba) was observed in cytosolic puncta on starvation, Atg8-PE-in the adult fly fat continued to be nuclear on starvation (Figure 4Bb). Specifically, we noted that the GFP::Atg8-PE-nuclear signal was the same between fed and starved states (Figure 4B’; p=0.98). Hence, when Atg8 is lipidation defective, Atg8 is nuclear on both fed and starved states.

Since Atg8-PE-displays a high level of nuclear localization in adult fly fat irrespective of the nutrient state (figure 4B), we predicted that Atg8-PE-would uncouple Upd2’s exogenous expression nutrient-responsive nuclear accumulation in fly fat, as it does in S2R+ cells (Figure 3A). To test this prediction, when we acutely expressed GFP::Atg8-WT or GFP::Atg8-PE-in adult fly fat (*ppl-gal4>TubGal80ts>UAS-Atg8-WT or UAS-Atg8-PE-*), and then assessed what happens to endogenous Upd2’s nuclear localization on starvation [*upd2-HA (genomic knock-in); ppl-Gal4tubGal80ts>UAS-Atg8-WT or Atg8-PE-*]. When Atg8-WT is acutely expressed, endogenous Upd2’s nuclear levels increase starvation within 4 hours (Figure 4C; p=1.0e-09). However, when we express Atg8-PE-acutely, the amount of nuclear Upd2 increases overall in the fed state compared to Atg8-WT, but nuclear Upd2 decreases on starvation (Figure 4C; p=0.003). Hence, exogenous lipidation defective Atg8 dysregulates Upd2’s nuclear levels from the systemic nutrient state *in vivo*.

Atg8-PE-expression prevented Upd2 nuclear accumulation (Figure 4C), and we wondered whether this impacted organismal starvation resilience. For conducting whole animal physiology experiments, we could not recover any viable progeny when we expressed Atg8-PE-in an Atg8 mutant background (Figure S5A). Since the expression of the Atg8-WT and Atg8-PE-transgenes were comparable on starvation (Figure S5B), we took the approach of over-expressing Atg8-WT or Atg8-PE-in fly fat and assessing systemic effects (See Study Design section below). First, we tested the ability of flies to mobilize fat stores and noted that flies expressing Atg8-WT (*Lpp-Gal4> UAS-Atg8-WT*) were able to break down their fat reserves on starvation (Figure 4Da; 50% mobilization; p=0.04). But flies with an exogenous expression of lipidation-defective Atg8-PE-in fly fat (*Lpp-Gal4> UAS-Atg8-PE-*) were unable to mobilize their TAG stores on starvation (Figure 4Da); this correlated with the reduced starvation resilience of fat specific expression of Atg8-PE-(Figure 4Ea).

Next, we went one step further and predicted that the inability of flies with fat-specific Atg8-PE-over-expression to mobilize TAG on starvation is at least in part due to Upd2’s cytosolic localization. If this prediction were to hold up, we should observe that the negative effects of Atg8-PE-over-expression are mitigated in the absence of Upd2. Consequently, we performed the TAG mobilization and starvation survival experiments in an *upd2-deletion* background (*upd2Δ; Lpp-Gal4> UAS-Atg8-PE-*). In line with our prediction, we found that exogenous expression of Atg8-PE-in fly fat of *upd2Δ* flies led to TAG mobilization (Figure 4Db) and starvation resilience (Figure 4Eb) at levels comparable to Atg8-WT transgene expression. In sum, these results support a model that Atg8’s lipidation, in addition to its critical role in autophagy, is required for Upd2 retention; this, in turn, determines how an organism survives prolonged starvation.

### UPD2’S NUCLEAR RETENTION IS CRITICAL FOR STARVATION-INDUCED FAT MOBILIZATION AND HUNGER-DRIVEN FEEDING

To further assess what happens when the inverse correlation between nutrient state and adipokine nuclear accumulation is genetically perturbed. We subjected flies expressing control transgene, Upd2-WT or Upd2-AIM-specifically in fly fat cells, to a high sugar diet (HSD) regime (See Methods and Study Design sections). We found that over-expression of Upd2-AIM-renders flies sensitive to an HSD regime (Figure 5A), suggesting holding Upd2 in the nucleus during a surplus diet where Upd2 is less nuclear (Figure S6A) reduces HSD survival. Conversely, we predicted that genetically ‘holding’ Upd2 in the nucleus during starvation, a state where endogenous Upd2’s nuclear accumulation increases (Figure 1A), will allow flies to survive starvation. Consistent with our prediction, the over-expression of Upd2-AIM-in fly fat cells (*upd2Δ; Lpp-Gal4> UAS-Upd2-AIM-*) on starvation provided resilience; Upd2-AIM-flies survived significantly longer than control or Upd2-WT overexpression (Figure 5B). Overall, this suggested that the inverse correlation between adipokine nuclear accumulation and nutrient state is important.

**Figure 5.**
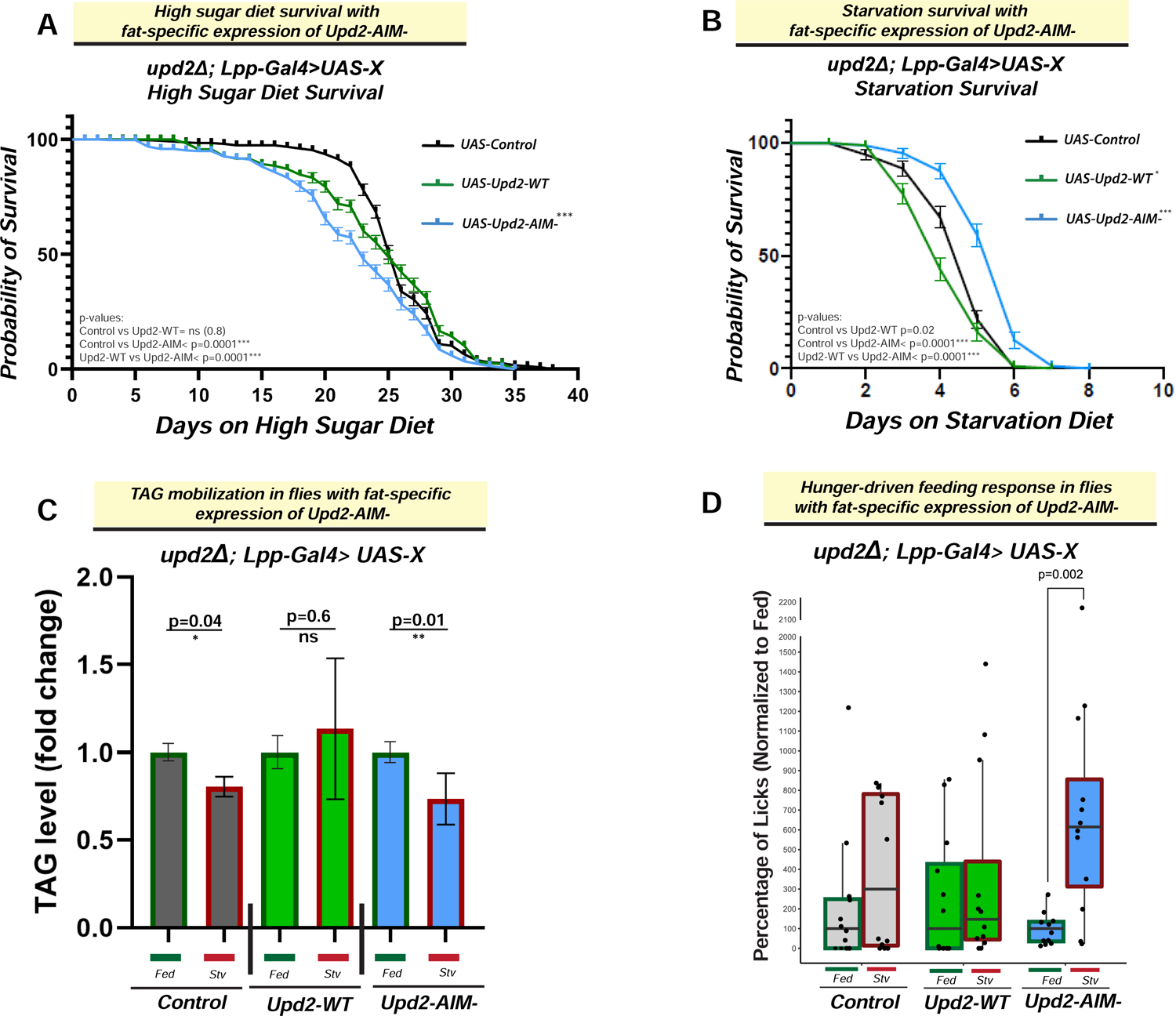
Upd2’s nuclear retention promotes starvation resilience and hunger-driven feeding. (A) and (B) <u>Effect of fat-specific expression of Upd2-AIM- on high sugar diet and starvation diets:</u> Flies expressing cDNAs (UAS-X) for Upd2-WT, Upd2-AIM- and control (UAS-Luciferase) specifically in fly fat (*upd2Δ Lpp-Gal4>UAS-X*) are subjected to (A) 30% high-sugar diet (HSD) and (B) 1% sucrose agarose diet. Per genotype >100 flies were used per experiment. Data shown is a consolidation of three independent experiments. Statistical differences in starvation survival were quantified using the Log-rank (Mantel-Cox) test. See also companion Figure S6B for data on Upd2-NES^+^AIM^-^ starvation survival. (C) <u>Effect of fat-specific expression of Upd2-AIM-on triglyceride mobilization during starvation:</u> Flies expressing cDNAs (UAS-X) for Upd2-WT, Upd2-AIM- and control (UAS-Luciferase) specifically in fat of flies in a Upd2 deletion background (*upd2Δ; Lpp-Gal4>UAS-X.* TAG levels at fed state were held at 1, and fold change in TAG at starvation was calculated. Error bars represent standard deviation of 9 biological replicates. Statistical significance is calculated using the two-tailed student’s T test. Statistical significance is calculated using the two-tailed student’s T-test. (D) <u>Effect of fat-specific expression of Upd2-AIM- on hunger-driven response:</u> Graph depicts boxplots of lick events of 7d old adult flies overexpressing cDNAs (UAS-X) for Upd2-WT, Upd2-AIM- and control (UAS-Luciferase) specifically in fly fat (*upd2Δ; Lpp-Gal4>UAS-X*) assessed for feeding behavior using FLIC (Fly liquid interaction counter- see methods). Flies experience either uninterrupted access to food (Fed *ad libitum*) or a 16 hour fast on 1% agarose (Stv). Number of licks was determined through contact using a fly liquid interaction counter (FLIC) on fed and stv flies in parallel for the first 3 hours after the 16-hour starvation period of the Stv group flies. Since flies were monitored in parallel, data was normalized as a percentage to the median fed lick values for each respective genotype, with median fed values being considered 100% lick events. Black dots represent individual fly licks (n=12). Significant differences between fed and starved groups were calculated using a t-test. Note only the UAS-Upd2AIM show a significant increase in licks events in the starved group compared to the fed control (p=0.002).

We wondered whether retention of Upd2 within the cell, but not specifically in the nucleus, would be sufficient to have the same systemic effects on starvation resilience (Figure 5B). To tease apart this nuclear versus cytosolic retention, we examined a state in which Upd2 was withheld in the cytosol but not released. To achieve this, we ectopically localized Upd2-AIM- to the cytosol by appending a constitutive nuclear export signal (NES) to Upd2-AIM- (See Methods). Expressing cytosolic Upd2-AIM- in the fly fat (*upd2Δ; Lpp-Gal4>Upd2-NES+-AIM-*), as predicted, was sufficient to re-localize Upd2-AIM- to the cytosol (Figure S6B; See arrows), whereas Upd2-AIM-expressing flies displayed no cytosolic localization (Figure S6B & Figure 2). Consistent with the role of Atg8 in mediating Upd2’s secretion, we found that, despite being localized to the cytosol (Figure S6D), Upd2-NES+-AIM- was not secreted (Figure S6E). Significantly, however, we noted that fat-specific expression of Upd2-NES+-AIM-(*upd2Δ; Lpp-Gal4>Upd2-NES+-AIM-*) did much worse on starvation than control flies (Figure S6C). These results suggested that it is *<u>not just retention</u> <u>within the cell, but specifically Upd2’s retention in the nucleus</u>* critical for exerting the systemic effects on starvation resilience.

We had previously shown that overexpression of Upd2-WT transgene in fly fat renders it incapable of mobilizing TAG reserves on starvation (Figure 2G in Rajan and Perrimon, 2012). Hence, we wanted to assay TAG mobilization in flies over-expressing Upd2-WT versus Upd2-AIM-specifically in fly fat (*upd2Δ; Lpp-Gal4> UAS-Upd2-WT/ AIM-*). We found that while Upd2-WT flies could not mobilize TAG on starvation (Figure 5C; Upd2-WT Fed vs. Stv p=0.6), Upd2-AIM-flies could mobilize TAG stores (Figure 5C; Upd2-AIM-Fed vs. Stv; p=0.01). Notably, the increased TAG mobilization of fat-specific Upd2-AIM-expressing flies concurred with increased starvation resilience of Upd2-AIM-expressing flies (Figure 5B).

Next, we assayed whether Upd2-AIM-displayed different feeding behavior post-starvation than Upd2-WT over-expression flies. Expressing Upd2-WT transgene, specifically in fly fat (*upd2Δ; Lpp-Gal4> UAS-Upd2-WT*), displayed no significant increase in hunger-driven feeding, as measured by the FLIC assay (Figure 5D, also see methods); this is consistent with Upd2’s role as a satiety signal. Conversely, Upd2-AIM-fat-specific over-expression flies (*upd2Δ; Lpp-Gal4> UAS-Upd2-AIM-*) displayed a significant increase in hunger-driven feeding (Figure 5D; Upd2-AIM-Fed vs. Starved; p=0.002). These findings suggest that nuclear accumulation of Upd2 on starvation exerts systemic effects; it promotes starvation resilience, fat-mobilization, and hunger-driven feeding.

### UPD2’S NUCLEAR RETENTION DURING STARVATION ALTERS GENE EXPRESSION

Our experiments reveal that nuclear retention is critical for Upd2-AIM- to exert systemic effects on starvation resilience. We hypothesized that Upd2 nuclear retention, during starvation, has some function within the nucleus. We wondered whether one such function might be for nuclear Upd2 to participate in gene regulation (See Discussion). To this end, we surveyed the bulk transcriptome of flies control flies (*Lpp-Gal4*), *upd2-deletion* flies (*upd2Δ; Lpp-Gal4*), and flies with fat-specific Upd2-AIM-over-expression (*upd2Δ; Lpp-Gal4>UAS-Upd2-AIM-*) and Upd2-WT over-expression (*upd2Δ; Lpp-Gal4>UAS-Upd2-AIM-*). We then analyzed differentially expressed (DE) genes between fed and starved states for each genotype mentioned above (Figure 6, S7).

**Figure 6.**
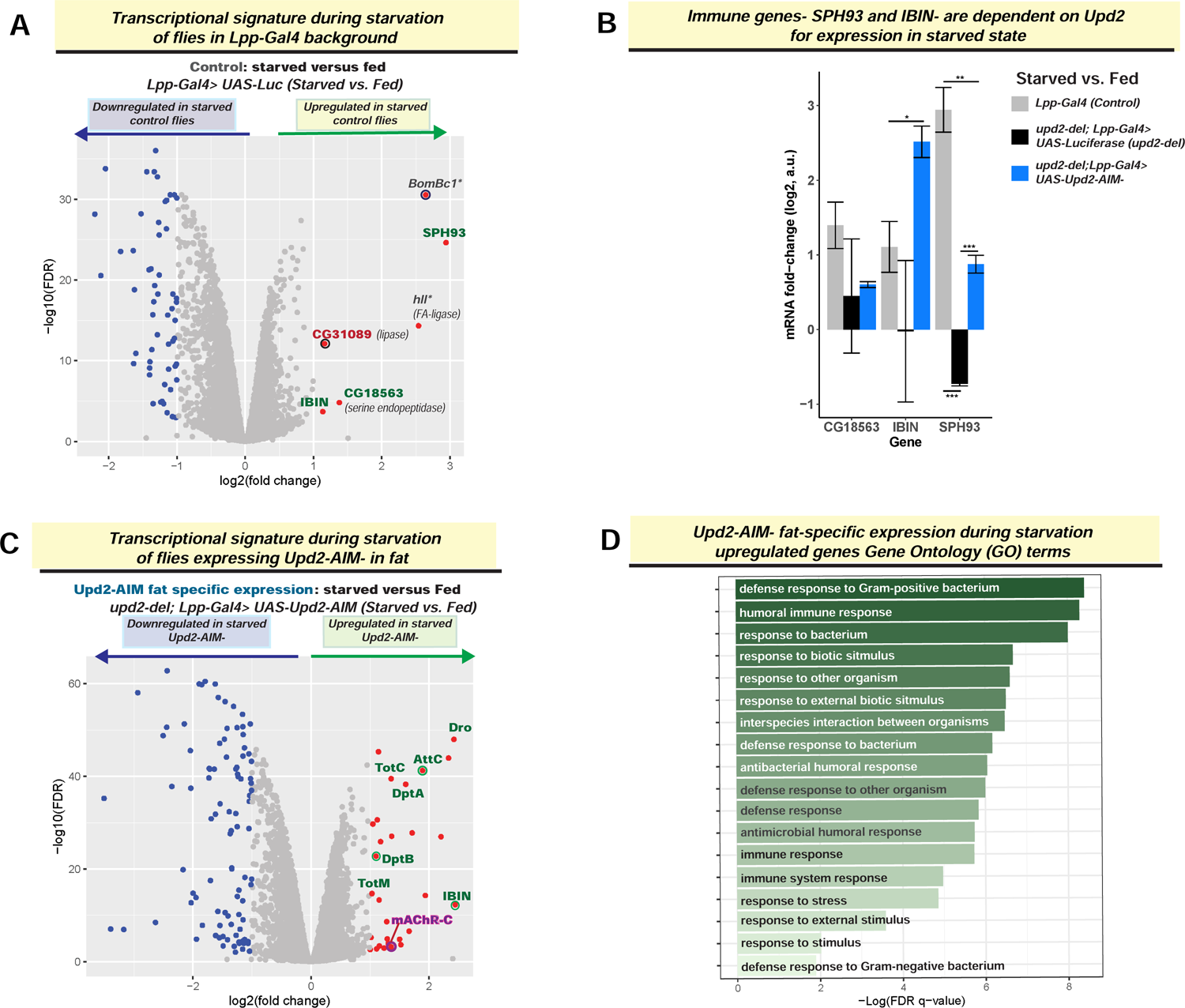
Upd2’s nuclear retention during starvation alters gene regulation. (A) and (C) <u>RNA-seq transcriptomic analyses of control and Upd2-AIM-over-expressing flies:</u> Volcano plot depicts starvation-induced changes in transcriptome, as assayed by RNA-seq on whole flies in control (A; *Lpp-Gal4> UAS-Luciferase*) and Upd2-AIM-fat specific overexpression (C; *upd2Δ; Lpp-Gal4> UAS-Upd2-AIM-*). (B) <u>Genes which are upregulated on starvation only in the presence of Upd2:</u> The immune genes - SPH93 and IBIN - are only upregulated on starvation in the presence of Upd2. The bar graph depicts the mRNA log2 fold change in response to starvation of genes in Y-axis. Each bar represents difference in fold change between starved and fed state of the indicated genotype. Error bars represent the standard deviation of log2 fold change among biological replicates (n=3). Significance was calculated using t-test and followed by *post-hoc* analyses of q-values to account for false discovery rate. Values of q<0.05 were considered significant. (D) The GO biological processes of secreted immune response genes are significantly upregulated in starved flies expressing Upd2-AIM-.

Systemic effects of starvation-induced Upd2 nuclear retention are likely to arise from increased and decreased gene expression. But we focused our analyses on upregulated genes since there were only six in controls on starvation (Figure 6A). Four of the six upregulated genes were involved in innate immune processes (SPH93, IBIN, Cg18563, BomBc1), and the other two highly upregulated genes were lipases (Figure 6A). Among the four genes with GeneOntology (GO) classification of innate immunity, two of them, including a secreted immune protein SPH93 (Serine protease homolog 93) and IBIN (Induced by Infection), relied on the presence of endogenous Upd2 for their starvation-induced expression (Figure 6B). Significantly, these two genes – IBIN and SPH93 – were highly upregulated in Upd2-AIM-over-expressing flies (Figure 6B). Overall, this suggested that nuclear Upd2 localization on starvation is required to express certain immune genes that encode secreted products (See Discussion).

We then performed deeper analyses for all the upregulated genes in Upd2-AIM-fat-specific expressing flies on starvation (*upd2Δ; Lpp-Gal4> UAS-Upd2-AIM-*; Figure 6C). We noted that in addition to one gene (mAchR-C) implicated in promoting feeding in mammals [63], most of the genes that were upregulated included secreted antimicrobial peptides (Figure 6C). When we analyzed the GO terms for genes upregulated in Upd2-AIM-during starvation, the key GO terms (Figure 6D) were all related to defense response to the bacterium. Hence, the over-expression of Upd2-AIM-during starvation upregulates a secreted immune peptide gene signature that provides a potential explanation for how it exerts systemic effects (See Discussion). Notably, overexpression of Upd2-WT transgene, which differs from AIM only in two amino acid sequences, and is expressed at relative levels to Upd2-AIM-transgene in baseline (Figure S7Ba), and starved states (Figure S7Bb), does not result in upregulation of this immune GO signature (Figure S7Ab). Hence, this supports the idea that increased nuclear retention of Upd2 on starvation promotes starvation resilience by upregulating a secreted innate immune peptide signature. In future work, directed experiments will be required to ask whether the upregulation of secreted immune genes contributes to the increased TAG mobilization, feeding behavior, and starvation resilience observed in Upd2-AIM-(See Discussion).

In conclusion, our results suggest that we have revealed a specific intersection node between cell-intrinsic and extrinsic mechanisms in regulating an organism’s response to nutrient extremes. We demonstrate that Atg8 regulates nutrient state-dependent localization of the *Drosophila* Leptin, Upd2 in both fed and starved cells (Summary of findings in Figure 7). Upd2’s nuclear accumulation on starvation, as a direct result of Atg8’s lipidation, is important for organismal adaptation to nutrient deprivation. Thus, we propose that Atg8’s role in the starvation response is not just limited to autophagy but also is required for adipokine retention. This process is critical for an organism to sense and adapt to starvation.

**Figure 7.**
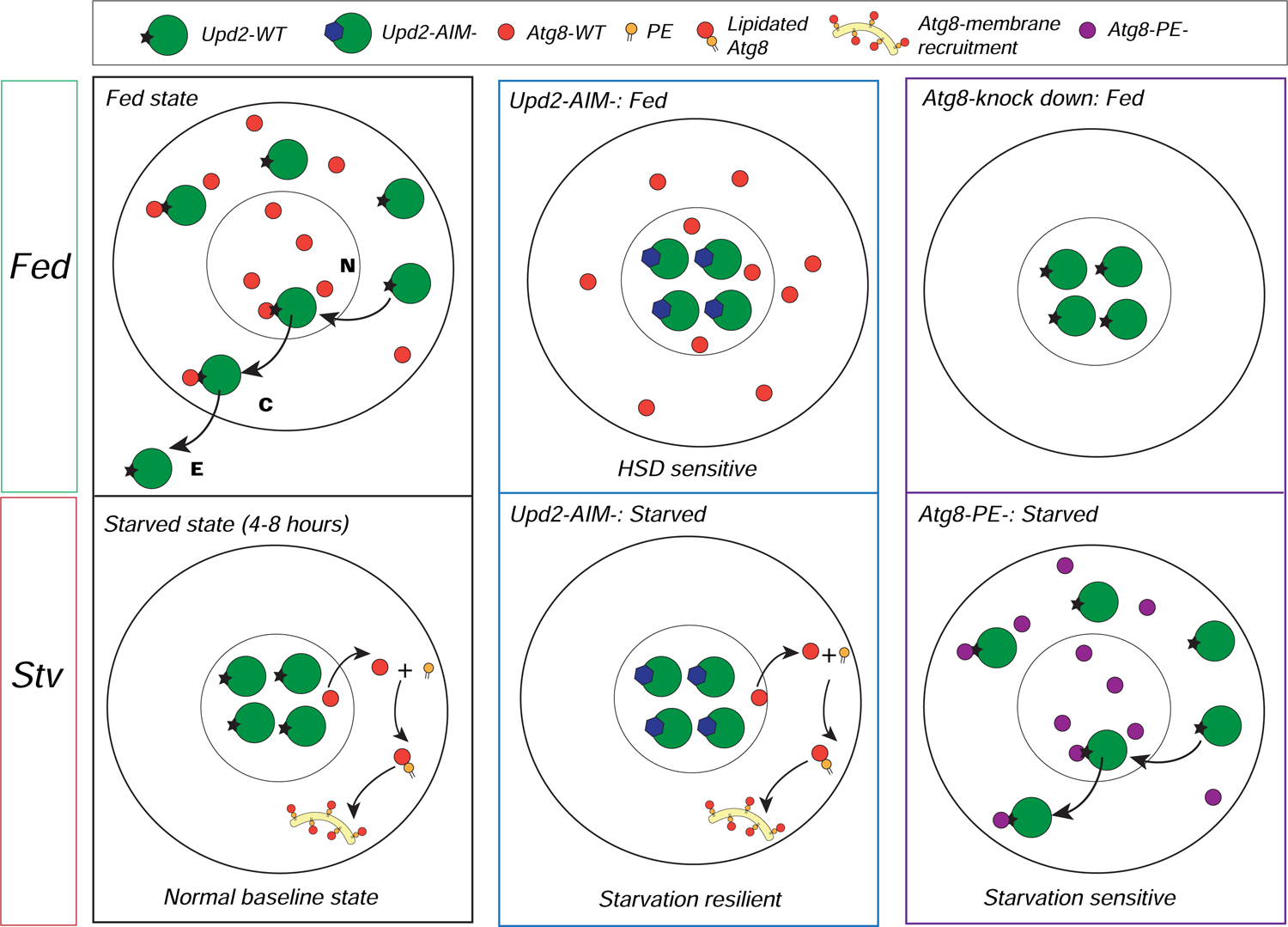
Summary of findings: Upd2’s intracellular localization is contingent on nutrient state and Atg8. Each illustration summarizes the key findings of the study. The upper panel is the key. Top three panels are fed state and lower three panels are in acute starvation (stv). Each panel illustrates a single cell. Notations: N-nucleus, C-cytosol, E-extracellular space.

- *Top left:* In a normal baseline fed state, the ortholog of human Leptin in flies, Upd2, enters the nucleus. Subsequently, Upd2’s Atg8-interaction motif (AIM) mediates interactions with Atg8 that enables Upd2 nuclear exit to the cytosol and then Upd2 is released to the extracellular space.
- *Bottom left:* During starvation, Atg8 transits to the cytosol and is lipidated. Depletion of the nuclear pool of Atg8 on starvation increases Upd2 nuclear accumulation and causes cells to retain the adipokine on starvation. Thus, Atg8’s lipidation state serves as a switch to license adipokine Upd2’s cytosolic or nuclear localization.
- *Top middle and right:* When Atg8’s interaction with Upd2 is disrupted (Upd2-AIM-) and when Atg8 levels are reduced by RNAi-mediated knock-down, Upd2 accumulates in the nucleus even in fed cells. This provides evidence for Atg8’s role in facilitating Upd2’s cytosolic localization in fed cells. We note that constitutive expression of Upd2-AIM- in fly fat renders flies sensitive to a HSD. This suggests that maintaining Upd2 in the nucleus, during a nutrient state where it is usually cytosolic, has a negative impact on resilience to a surplus diet.
- *Bottom middle:* Upd2-AIM-remains in the nucleus on starvation, while Atg8 translocates to the cytosol. Upd2-AIM-flies are starvation resilient and upregulate a systemic innate immune gene expression signature.
- *Bottom right:* Lipidation-defective Atg8-PE-continues to be nuclear on starvation. Presence of Atg8-PE-allows Upd2 to be cytosolic even on starvation and renders flies sensitive to starvation.

## DISCUSSION

### A NOVEL NON-CANONICAL ROLE FOR ATG8 IN NUTRIENT-STATE DEPENDENT PROTEIN LOCALIZATION

Upd2 is a 47 kDa protein. Given the structural restraint of nuclear pore complexes (NPC) to only permit proteins <40kDA to pass through [64], Upd2 is likely to require active nucleocytoplasmic (NC) transport mechanisms that enable transit through the NPC [65]. Consistent with the role for classic nuclear export being required, we find that Embargoed/CRM1 is required for Upd2’s nuclear exit (Figure S1C). But, using the NETNES mapper tool [66] did not identify a canonical NES in Upd2. Hence, we hypothesized that other adapter proteins interacting with Upd2 might promote its nuclear export. Unexpectedly, we found that mutations to Upd2’s AIM or KD of Atg8 phenocopies knockdown of nuclear export protein Emb (Figure 2, S2). This suggested to us that Atg8 is required for Upd2’s cytosolic localization in fed cells. Consistent with this nanobody-based reconstitution of the physical interaction between Upd2-AIM-, that is only localized to the nucleus, and Atg8 is sufficient to re-localize Upd2 to the cytosol (Figure 2D, 3B, 3B’).

Our study does not yet resolve how Upd2 localizes to the nucleus in the first place. But, based on the observations that Upd2 is nuclear when Atg8 is knockdown and when its AIM is disrupted, we conclude that Atg8 does not regulate Upd2’s nuclear entry. Analysis of Upd2’s sequence using a bioinformatic tool-classic NLS (cNLS) mapper [67–69], predicted that Upd2 contains an NLS at residues 141-150 (GRVIKRKHLE) with a score of 10.5. According to the cNLS mapper, a GFP reporter protein fused NLS, with a score of 10, is exclusively localized to the nucleus. Hence, we reasoned that Upd2 is likely to be nuclear, but when Upd2 is ‘licensed’ by cofactors like Atg8, it will Upd2 transit to the cytosol.

We considered the alternate hypothesis that Atg8 cytosolically tethers Upd2 to prevent nuclear entry. But if that were the case, we would expect that Atg8, which is highly cytosolic on starvation, would tether Upd2 to the cytosol. Given that Atg8 nuclear depletion, within 4 hours of acute starvation (Figure 4A), coincides with increased Upd2 nuclear accumulation (Figure 1A), we favor the hypothesis that the nuclear pool of Atg8 in fed cells facilitate Upd2 nuclear exit.

Further work will be needed to resolve all the events leading to Upd2’s Atg8-mediated nuclear exit. We postulated that Atg8, a ubiquitin-like protein (Shpilka et al., 2011), might operate akin to other ubiquitin and ubiquitin-like proteins that have been shown to function in cellular localization [70–72]. Specifically, SUMO (small ubiquitin-like modifier) positions its target protein (RanGAP1) to the NPC to increase the efficiency of its translocation [73]. Similarly, Atg8 might position adipokine Upd2 to enable efficient export via the classical nuclear export pathway or may serve to direct Upd2 to other nuclear export mechanisms such as nuclear budding [74].

How widespread this role for Atg8 in nucleocytoplasmic protein localization remains resolved. But it is quite plausible that other proteins, such as ribophagy receptor NIUFIP1 [75] with 4 AIM sequences, and localizes to cytosol on starvation, might utilize Atg8 for context-dependent NC shuttling. Furthermore, insulin retention on acute starvation is a critical response in flies and mammals [76]. It will be important to explore whether Atg8 mediated retention, like Leptin/Upd2 in fat, also applies insulin in pancreatic beta cells. In sum, Atg8’s controls context-dependent nucleocytoplasmic localization could be another layer of regulation in controlling protein localization during starvation and merits further investigation.

Atg8 controls the two step-process for Upd2 release in fed cells – 1: nuclear exit and 2: GRASP-mediated extracellular-release (Figure S2D, S2E). This study shows that lipidation-defective Atg8 (VHH-Myc::Atg8-PE-) can localize Upd2-AIM- to the cytosol (Figure 3Bd), but we do not know whether the lipidation defective Atg8 can restore Upd2-AIM-‘s secretion defect. Debnath and colleagues have comprehensively delineated how Atg8/LC3 lipidation is critical to Atg8’s role in extracellular vesicle secretion, a process that they term LDELS [40]. Based on their elegant study [40], we propose that though Upd2 can transit to the cytosol in the presence of lipidation-defective Atg8, it is not secreted. Further studies will be needed to resolve whether Upd2 undergoes an LDELS secretion process in *Drosophila* fat cells.

### HOW DOES THE NUCLEAR RETENTION OF UPD2 EXERT SYSTEMIC EFFECTS ON STARVATION?

In this work, we reveal that the adipokine Upd2 needs to be retained within the nucleus for exerting its positive effect on starvation resilience (Figure S6B, C). Our bulk transcriptomic analysis hints at some possibilities for why Upd2-AIM-might be able to exert this systemic influence. First, we find very few genes are upregulated in overnight 16-hour starvation in our control background. Of them, two innate immune genes are dependent on upd2 for expression since they are not upregulated in a *upd2Δ* background. Significantly SPH93 is secreted in the extracellular space and required for gram-positive bacteria response [77–79]. IBIN, which encodes a long non-coding RNA (lncRNA) induced upon infection, also required Upd2 to express starved states (Figure 6B). IBIN enhances the expression of genes required for glucose retrieval [80]. Hence IBIN’s upregulation downstream of Upd2 could play in starvation resilience by increasing sugar utilization. Furthermore, in a background where Upd2 is highly nuclear (Upd2-AIM-), the GO signature of upregulated genes is that of secreted innate immune peptides (Figure 6D). Notably, Turnadot (Tot) family genes have been shown to provide humoral stress resistance from both infection and sleep deprivation [81, 82] and are upregulated downstream of JAK/STAT signaling [83]. Drosocin (Drs), which is highly upregulated in the presence of Upd2-AIM- on starvation (Figure 6B), promotes increased feeding motivation [84]. Similar DptB, another AMP over-expressed in Upd2-AIM-(Figure 6B), acts on the central brain circuits to alter long-term memory [85]. Hence, we postulate that increased AMP expression in Upd2-AIM-may impinge on systemic processes, perhaps by altering brain function, and this might underlie their increased hunger feeding motivation (Figure 5D). Furthermore, the increased lipolysis we observe with Upd2-AIM over-expression flies (Figure 5C) aligns with the emerging role in mammalian studies for AMPs in TAG mobilization [86, 87]. In sum, our current working hypothesis is that the secreted immune peptide signature downstream of Upd2 nuclear retention enables the organism to adapt to starvation and promotes hunger-driven feeding. Though it is beyond the scope of the current study, testing these specific hypotheses will be refined and developed in future work.

### HOW COULD UPD2 AFFECT GENE EXPRESSION?

Transcriptional and epigenetic regulation of feeding behavior [88] and glucose sensing [89–91] is now appreciated. But how does Upd2, which does not have a DNA-binding sequence, participate in gene regulation? Upd2, like leptin, a ligand of the JAK/STAT signaling cascade [45], leads to the activation of the STAT transcription factor. Whether nuclear retention of Upd2 activates transcription downstream of STAT in an ‘autocrine’ fashion; or whether Upd2 fine-tunes gene expression by binding to other DNA and chromatin binding factors will need to be explored in future work. The latter hypothesis is supported by preliminary findings of our lab’s proteomic datasets. In multiple independent proteomic surveys of Upd2 immunocomplexes, we identified ten bonafide DNA binding and chromatin binding factors that complexed with Upd2 (Kelly and Rajan, unpublished results); whether these are bona fide interactors and if they have a functional significance will be tested in future work.

An intriguing observation is that Upd2-AIM-upregulates the innate immune gene signature only on starvation. Why does Upd2-AIM-not upregulate starvation responsive genes even in the fed state? One possibility is that factors required for nuclear Upd2 to engage in gene expression become available only on starvation. Another possibility is that nuclear factors in the fed state might prevent Upd2-AIM-from accessing its gene expression partners. Could Atg8 itself be such a factor that prevents Upd2 from engaging in gene regulation during the fed state? Atg8 signal in the nucleus is directly correlated with the nutrient state. It is upregulated on an HSD regime and significantly reduced in fat nuclei within 4 hours of starvation (Figure 4A). Hence, we speculate that Atg8, though it cannot bind Upd2-AIM, could sequester factors required for Upd2-AIM’s access to gene regulatory function in fed cells. This hypothesis is consistent with recent reports that Atg8 can regulate gene expression in *Drosophila* fat by working with transcription factors [92]. Even if we discount the Upd2-AIM-effects on gene expression as a gain-of-function effect, we note that Upd2 is required to upregulate gene expression at endogenous levels in starved flies (Figure 6B). Upd2 is likely to be nuclear localized in the endogenous expression state because of a strong NLS. Therefore, an active mechanism to prevent improper gene regulation by Upd2 in fed cells is likely to be in place. Hence, Atg8 itself, in addition to promoting Upd2’s nuclear exit (Figure 2), might prevent improper gene regulation by Upd2. In sum, our study has generated many testable hypotheses regarding Upd2’s potential role in gene regulation. We hope that this will stimulate future work.

### WHAT IMPLICATIONS DO DROSOPHILA UPD2 STUDIES HOLD FOR MAMMALIAN LEPTIN BIOLOGY?

Leptin does have two putative AIM-like sequences. But unlike Upd2 (47kDa), leptin (15 kDa) is unlikely to require facilitated nuclear export. Even if Atg8-mediated nucleocytoplasmic shuttling does not apply to mammalian leptin, it is possible that, like Upd2, Leptin utilizes a secretory pathway mediated by Atg8. This is not a far-fetched possibility given that human leptin, just like Upd2, requires unconventional secretion [27]. Hyperleptinemia, i.e., high levels of circulating leptin, has been identified as a primary cause of ‘leptin resistance’- a state wherein leptin cannot effectively signal satiety state to the brain [93]. This resulted in dysfunctional feeding behavior and decreased energy expenditure. Thus, our report that Atg8’s lipidation retains *Drosophila* Leptin/Upd2 represents a novel avenue to develop interventions to improve Leptin sensitivity.

This study uncovered a possible link between adipokine nuclear retention on starvation and the upregulation of antimicrobial peptides (AMPs). A recent study showed that in mice, gut cells upregulate AMP production. It was suggested that increased AMP production in non-feeding states is an anticipatory mechanism to prevent infection. Significantly, Hooper and colleagues’ study found that the increase in gut AMP production is controlled by the transcription factor STAT3 [94]. Strikingly, leptin also activates the STAT3 transcription cascade [95]. Hence, we suggest that, like Upd2, Leptin retention in adipocytes could increase immunity by promoting AMP expression downstream of STAT3.

In conclusion, the study on how *Drosophila* Leptin-Upd2 – is acutely retained in fly fat cells on starvation has led to numerous unanticipated insights raises many questions that should stimulate work both in invertebrate and mammalian systems.

### STUDY DESIGN-LIMITATIONS AND CONSIDERATIONS

*Atg8:* In *Drosophila,* clonal analysis of larval fat tissues is the standard for studying how autophagy (Atg) core pathway genes affect cell-intrinsic processes [96]. While clonal analysis addresses is an elegant genetic tool to study cell-intrinsic functions of Atg genes, it is not compatible with experimental design to study whole animal physiology. Therefore, for the physiology assays, in which we assess how Atg8’s lipidation in fat affects TAG mobilization and starvation resilience (Figure 4D, 4E), we have compared the effect of Atg8-WT transgenic over-expression with the Atg8-PE-in fly fat. The reason for this design is two-fold. The first is to overcome the hurdle of studying a specific function of Atg8, its lipidation, which is critical to animal survival. We note that lipidation-defective Atg8 flies do not survive (Figure S5A). Hence the only viable strategy to study the effect of defective Atg8 lipidation is to utilize fat tissue-specific expression of Atg8-PE- and compare its effects to Atg8-WT. It is important to note that we compare the effects of two transgenes-Atg8-PE- and Atg8-WT-head-to-head. These transgenes differ in a single amino acid (the penultimate glycine of Atg8) and are expressed at comparable levels even on starvation (Figure S5B).

Furthermore, these results will be viewed in conjunction with the V_H_H-nanobody based reconstitution of interactions between Atg8-PE- and Upd2, where Atg8-PE-increases Upd’2 cytosolic exit, even on starvation, despite the presence of endogenous Atg8 (Figure 3A). Secondly, our goal is to uncover the systemic effects of only altering Atg8’s lipidation status in the fat. Since Atg8 has roles in every tissue, the most feasible design is to use the tissue-specific expression of lipidation-defective Atg8 and assess how it impacts whole organism physiology in relation to Atg8-WT. Hence, within the limitations of the system, we have utilized internally controlled experimental approaches to assess the effect of Atg8’s lipidation status on endogenous Upd2 localization and whole animal physiology.

For experiments to study the effect of nutrition on protein-localization (Figure 4A and B), we utilized the UAS/Gal4 system for specifically expressing Atg8-WT and Atg8-PE-transgenes in the fly fat. This design of driving protein expression downstream of fat-specific Gal4 allows us to uncouple transcription and translation-dependent effects of nutrient state on Atg8 mRNA and protein levels. Hence, we quantify the nuclear GFP signal in fat cells in 3D-volume, representing Atg8 levels. Furthermore, we note that on starvation, both Atg8-WT and Atg8-PE-transgenes show a similar fold change in gene expression (Figure S5B). Nonetheless, Atg8-WT is cytosolic on starvation (Figure 4A), but Atg8-PE- is nuclear (figure 4B). This lends credence to our claim that we follow the nutrient state effect on protein localization rather than gene expression.

*Upd2:* Upd2 is the functional fly ortholog of human leptin [19] and plays a fat-specific role in regulating the systemic nutrient state. But, as shown by multiple independent groups, Upd2 exerts systemic effects by signaling from other organs and, specifically, coupling gut metabolic state to the olfactory system[97] or acting in hemocytes to regulate immunity [98]. We have utilized fat-tissue-specific drivers to distinguish Upd2’s adipokine-specific function from its other organ-specific roles. *In lieu* of the ability to perform longitudinal diet-related experiments (10-40 days) with a knockdown of Atg8 in fat, which is lethal over seven days, we utilized Upd2-AIM-over-expression in fly fat. Upd2-AIM-phenocopies Atg8-KD (Figure 2A, 2B). Using this design, we have assessed the systemic effects of various Upd2 transgenes (*Upd2-WT, Upd2-NES+-AIM-and Upd2-AIM-*).

Given that these transgenes are all inserted in the same genome site (attP40) and have similar expression levels (Figure S7B), it allows for interpretation of the specific mutation to Upd2’s Atg8 binding domain means. One valid reservation is that we reveal are gain-of-function effects of Upd2-AIM-. Even if that were to be the case, it is striking that we don’t see these gain-of-function effects on starvation resilience when we retain Upd2-AIM-in the cytosol (*Upd2-NES+-AIM-)* (Figure S6A, S6B); this suggests that retention of Upd2 in the nucleus on starvation is biologically meaningful. Hence, within the limitations of the system, we have utilized internally controlled experimental approaches to assess the effect of Upd2’s nuclear localization on whole animal physiology.

## ACKNOWLEDGMENTS

We are grateful to the late Suzanne Eaton for the Lpp-Gal4 flies. We thank Laura Holderbaum for assistance with cell culture and immunoprecipitation experiments. Zachary Goldberg for technical assistance with setting up the FLIC assay. We thank Alyssa D. Dawson and Dolores Covarrubias at the Fred Hutch Genomic Shared resources facility for assistance with RNAseq. We are grateful to Feinan Wu at the Fred Hutch Bioinformatics core for RNAseq analysis. We thank David Strutt for sharing sequences for designing the V_H_H tagged constructs. We thank Susan Parkhurst for her advice on the NES sequences and critical comments on the manuscript. Funding: This work is possible due to grants awarded to AR from NIGMS (GM124593) and New Development funds from Fred Hutch. KPK was supported by the NIH Chromosome Metabolism and Cancer Training Grant (T32CA009657) and is currently supported as an NSF Post-Doctoral Research Fellow (NSF Award #2109398). Genomic reagents from the DGRC, which is funded by NIH grant 2P40OD010949, were used in this study. Stocks obtained from the Bloomington Drosophila Stock Center (NIH P40OD018537) and Transgenic RNAi Resource project (NIGMS R01 GM084947 and NIGMS P41 GM132087) were used in this study. Author contributions: Conceptualization: AR, PR; Investigation: MEP, CES, AEB, KPK, AM, AR; Formal analysis, Data Curation: MEP, CES, AEB, KPK, AM, AR; Visualization: MEP, CES, AEB, KPK, AM, AR, JD; Supervision: AR; Writing-original draft: AR; Writing-review and editing: PR, KPK, AM. Funding acquisition: AR. Competing interests: Authors declare no competing interests.

## RESOURCE AVAILABILITY

### Lead Contact

Requests for further information, reagents, and resources should be directed to and fulfilled by the Lead Contact, Akhila Rajan.

### Materials Availability

*Drosophila* strains generated in this study are available from the corresponding author, Akhila Rajan.

### Data and Code Availability

The datasets generated in this study are available from the corresponding author, Akhila Rajan.

## EXPERIMENTAL MODEL AND SUBJECT DETAILS

### Experimental Animals

*Drosophila melanogaster* Only males were used in experiments at 5-10 days post-eclosion. Flies were cultured in a humidified incubator at 25°C with a 12h light-12h dark cycle and were fed a standard lab diet, containing per liter: 15 g yeast, 8.6 g soy flour, 63 g corn flour, 5g agar, 5g malt, 74 mL corn syrup. For acute starvation, adult male flies, seven days old, were subjected to 4-hour starvation on 0% sucrose in agar at 29°C. For survival curves, flies were subjected to 1% sucrose in an agar diet. For RNAi experiments, flies were raised at 18°C until seven days post-eclosion, after which they were shifted to 29°C for seven days.

The following fly strains used in this study were from our previous work [19, 27] and obtained from other investigators or the Bloomington *Drosophila* stock center (BDSC): upd2Δ3-62 (upd2Δ; [45]), *Lpp-Gal4* [99], Atg8-Trojan-Gal4 ([100]; BDSC 77836). The control UAS strain used for over-expression experiments is UAS-Luciferase [19]. RNAi lines from the TRiP facility at Harvard Medical School (http://www.flyrnai.org/TRiP-HOME.html) include: Luciferase-RNAi (JF01801), Atg8-RNAi #1 (JF02895, BDSC 28989), Atg8-RNAi #2 (HMS01328, BDSC 34340).

The following UAS lines were generated for this study: *UAS-Upd2::mCherry, UAS-Upd2-AIM-::mCherry, UAS-GFP::Atg8; UAS-GFP::Atg8-PE*-. All transgenes of the same gene were inserted at the same att site to control expression levels—either attp40 or attP2. For RNAi experiments and temporal over-expression in Figure 2B and Figure 4C, flies were generated with the following genotype: *Upd2-crGFP; Ppl-Gal4, TubGal80ts* was crossed to Atg8-RNAi, Luc-RNAi for 2B; For 4B and 4C and *UAS-GFP::Atg8-WT* and UAS-GFP::Atg8-PE-. The parents of this cross (F0 generation) were maintained at 18c for both the strains, F1s were allowed to eclose at 18c. A week after eclosion, the F1s were moved to 29c for 5-7 days before dissection and staining.

The ability of UAS-GFP::Atg8 and UAS-GFP::Atg8-PE-to rescue the loss of Atg8 function was tested by introducing Atg8 variant transgenes into the Atg8-Trojan-Gal4 line [100] in which insertion of the Trojan-Gal4 into the Atg8 locus generates a lethal allele. We confirmed the presence of Gal4 by crossing to UAS-GFP.

### Cell lines

*Drosophila* S2R+ cells were used for all cell culture-related experiments. This cell line was chosen because previous studies have validated their applicability to study autophagy 7-9 and protein secretion. The cells were maintained in Schneider’s medium (GIBCO), 10% heat-inactivated FBS (SIGMA), and 5% Pen-Strep (GIBCO) at 25°C. For starvation, cells were incubated in Schneider’s Insect Medium without Amino Acids (United States Biological, Cat# S0100-01.10) for the indicated amount of time.

## METHOD DETAILS

### Cloning and Transgenic Flies

All cloning was done using the Gateway® Technology. For Atg8, entry cDNA was PCR amplified from Atg8 cDNA (clone LD05816 DGRC-Gold collection) and cloned into pENTR-D/TOPO using BP reaction (Gateway® BP Clonase II enzyme mix, Cat#11789-020, Invitrogen). For *Drosophila* Upd2 variants, pENTR-Upd2 [27] made for previous work in our lab was used. For site-directed mutagenesis of putative AIM sites, pENTR-Upd2 was mutagenized to convert tryptophan and leucine encoding codons to alanine. For the addition of a canonical NES+, a sequence corresponding to amino acids LQELLELLRL [101] was inserted after the start codon in either pENTR-Upd2-WT or pENTR-Upd2-AIM-. For site-directed mutagenesis of the Atg8 lipidation site, pENTR-Atg8 was mutagenized to convert its penultimate Glycine to Alanine. All mutagenesis was done using the Q5® Site-Directed Mutagenesis Kit from NEB (Cat # E0554S). The sequence of oligonucleotides used for this mutagenesis reaction is available on request. The entry vectors were then moved using LR clonase reaction (Gateway® LR Clonase® II Enzyme mix, Cat#11791-020, Invitrogen) into destination vectors compatible with fly transformation, or cell culture with the appropriate C-terminal tags for Upd2 and N-terminal tags for Atg8. Transgenic flies were generated by using the microinjection service provided by Bestgene Inc.

For generating the V_H_H tagged constructs, the sequence for V_H_H tag and appropriate linkers, utilized in a prior study by Strutt and colleagues [57], was appended to either Myc (control: V_H_H-Myc), Atg8-WT (V_H_H-Myc::Atg8-WT), or Atg8-PE-(V_H_H-Myc::Atg8-PE-). Double-stranded sequences for the three constructs (available on request) were synthesized using Codex DNA Inc. Synthetic DNA platform and cloned into PDONR221 using BP clonase (Gateway™ BP Clonase™ II Enzyme mix; Catalog number: 11789020, Invitrogen). Then, to generate expression clones, the PDONR vectors were cloned into Expression competent S2 cell culture vectors from the Murphy Collection available from DGRC using LR clonase.

### Generation of C-terminal tagged Upd2 CRISPR lines

GFP or HA-tagged Upd2 CRISPR lines were developed in collaboration with WellGenetics Inc. using modified methods of Kondo and Ueda [102]. In brief, the following gRNA sequence was cloned into U6 promoter plasmids for HA lines: TCCAATGAGTCTTGAGCCCT[CGG]/GCCGAGGGCTCAAGACTCAT[TGG]. Cassette 3xHA RFP containing 3xHA and 3xP3 RFP and two homology arms were cloned into pUC57 Kan as donor template for repair. upd2/CG5988 targeting gRNAs and hs-Cas9 were supplied in DNA plasmids, together with donor plasmid for microinjection into embryos of control strain w1118. F1 flies carrying selection markers of 3xP3 RFP were further validated by genomic PCR and sequencing. CRISPR generates a break in upd2/CG5988 and is replaced by cassette 3xHA RFP. 3XP3 RFP, which facilitates the genetic screening, was flipped out by Cre recombinase. RE loxP RE (46 bp) remains after excision between Stop codon and 3’UTR. For GFP lines, gRNA sequence TCCAATGAGTCTTGAGCCCT[CGG] was used.

### Tissue culture

Before transfection, cells were passaged to 60-80% confluency. Cells were cultured in 96 well plates for transfections related to ELISA experiments and coimmunoprecipitations in 6-well plates. For ELISAs, cells were transfected with 20ng/well of pAc-upd2::GFP 3 or indicated Upd2 variants, 10ng/well pACRenilla::Luciferase, and 150ng of dsRNA/well for knockdown experiments. For coimmunoprecipitations, cells were transfected with 200ng/well of the indicated plasmid.

For imaging experiments, cells were seeded on poly-D-lysine coated 8-well chambered culture slides for fixed imaging (MatTek CCS-8). The T7 flasks, which were 100% confluent, were used for seeing cells at 1:10 dilution at 400μl per well of the 8-well chamber dish or 1:20 dilution at 200μl per dish for FluoroDish. Transfections were done with plasmids indicated. For 8-chamber slides, 100ng/well of each plasmid was transfected. Transfections were done using the Effectene kit (Cat# 301427, QIAGEN) per the manufacturer’s instructions.

### dsRNA production and cell treatments

Amplicons for dsRNAs were designed using the SnapDragon dsRNA design tool (https://www.flyrnai.org/snapdragon) and in vitro transcribed (IVT) using MEGAscript® T7 Transcription Kit (Cat# AMB1334-5, ThermoFisher). IVT reactions were carried out as per the protocol provided by the DRSC (available at: http://www.flyrnai.org/DRSC-PRS.html). The sequence of amplicons used in this study can be found in the table below. LacZ dsRNA was used as controls. All dsRNA knockdown experiments were carried out using two independent dsRNAs per gene. We found that this produced a knockdown efficiency of >85% (based on qPCR analysis) in S2R+ cells. S2R+ transfected with dsRNAs were incubated for four days to allow for gene knockdown. On the 4th day, the media was changed, and the ELISA assay was carried out on the 5th day. Note that the data is represented as percent change in Upd2/Leptin secretion normalized to transfection efficiency with 0% change indicating baseline secretion level. See the ELISA assay procedure below.

### Primer sequences used for dsRNA production

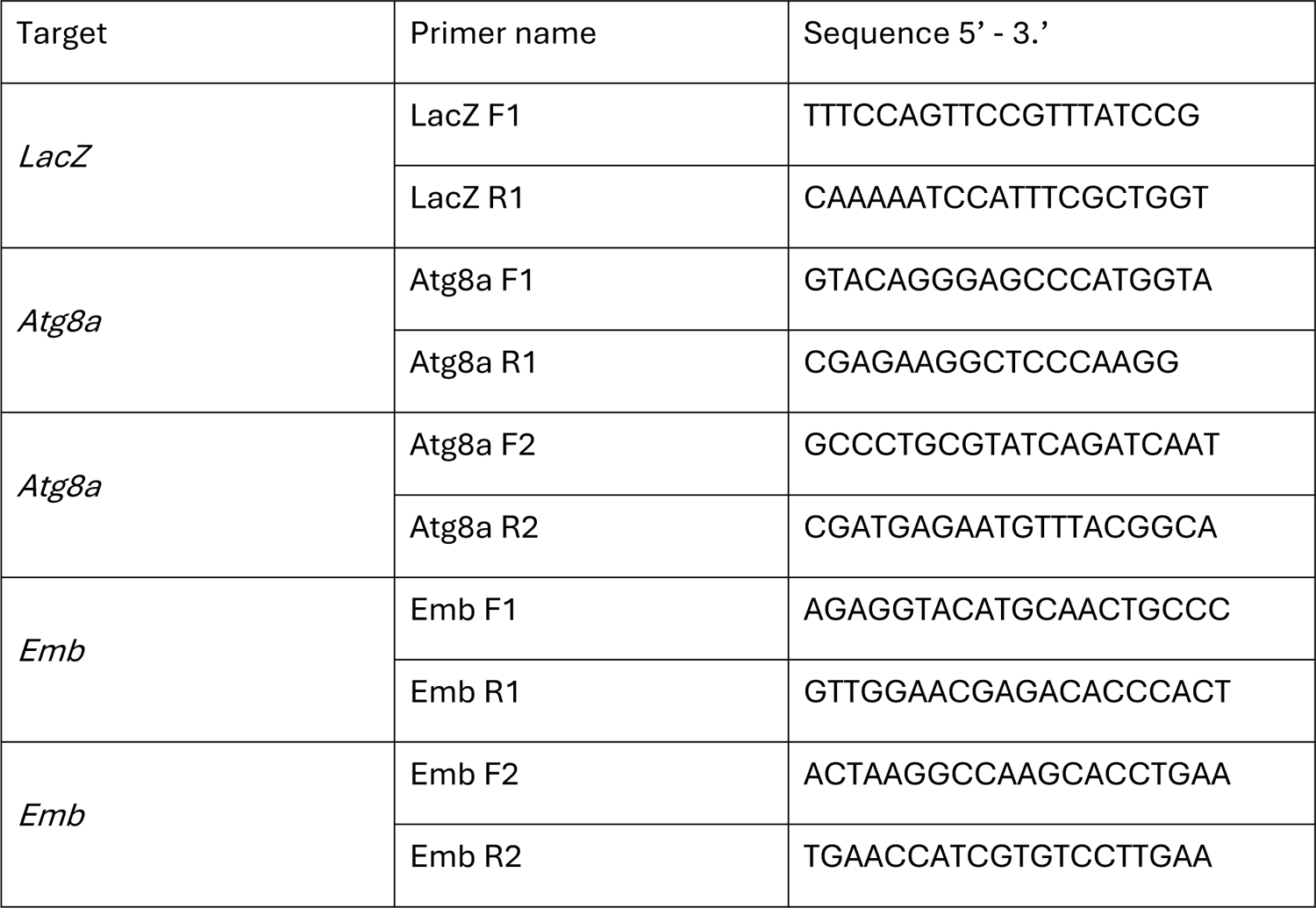

#### Treatment of cells with drugs

For drug treatment experiments, the media was replaced with media containing the drug on day 3 after transfection with upd2::GFP. 4 hours later; the conditioned media was used for ELISA or imaging. Drugs used in this study include Torin1 (Cat# ab218606, Abcam). Stocks solutions of the drugs were made in DMSO as per the manufacturer’s instructions and used at a working concentration indicated in the Figure S3 legends. DMSO treated replicates were used as controls. Note that for ELISAs, the data is represented as percent change in Upd2 secretion normalized to transfection efficiency, with 0% change indicating baseline secretion level.

#### ELISA assays

GFP sandwich ELISA assay was used for detecting Upd2::GFP. On day 1, 96 well medium bind polystyrene plates (Cat#CLS3368-100EA, Sigma) were incubated overnight at 4C with coating antibody (Cat# ACT-CM-GFPTRAP, Allele Biotechnology) diluted in 0.01M pH8.0 bicarbonate buffer at a concentration of 1μg/ml. On day 2, plates were washed briefly with PBS, blocked for 30 minutes with 1% BSA block in PBS, coated with conditioned media, and incubated overnight at 4C. Recombinant GFP protein (Cat# MB-0752, Vector labs), diluted in S2R+ cell growth media (64ng/μl to 4 ng/μl), was used in every ELISA plate as a positive control to ensure linearity of GFP readings. On day 3, the plates were washed with PBS+0.05% Tween-20 (PBS-T) blocked with 1% BSA in PBS for 30 minutes at RT. GFP detection antibody (Cat# 600-401-215, Rockland) was added to the diluted 1:1000 in 0.1% BSA in PBS-T. Plates were washed with PBS-T and incubated with secondary HRP conjugated anti Rabbit secondary antibodies (Cat# ab136636, Abcam) diluted at 1:5000 in 1% BSA block. Plates were washed in PBS-T with a final wash in PBS. For detection, each well was incubated 100μl 1-step Ultra-TMB ELISA substrate (Cat# 34028, Pierce), which was previously equilibrated to RT, for approximately 5-15 minutes until detectable blue colorimetric reaction occurred. The reaction was stopped with 2N sulphuric acid, and absorbance was measured at 450nm. The TMB readings were normalized to transfection efficiency as measured from Renilla Luciferase assays (see below).

#### Renilla Luciferase assay

On day 2 of the ELISA assay, after the conditioned medium was transferred for use in ELISA assays, cells were re-suspended in 50μl of PBS and incubated with 50μl/well Renilla-Glo® Luciferase reagent (Cat# E2710, Promega) for 10 minutes and read using a multiwell luminometer.

#### Quantification for ELISA assays

For ELISA assays, the ELISA signal readings are normalized to transfection efficiency; the data is represented as percent fold change from control used as the baseline. Specifically, the ratio of TMB readings to Renilla Luciferase readings is calculated. This ratio from the control is used as a baseline, and the data is represented as the percent fold change of experimental conditions with respect to the control. A 2-tailed t-test quantified statistical significance on 6 biological replicates per condition. Error bars represent %SD (Standard Deviation). For TAG analysis, statistical significance was quantified by a 2-tailed t-test on 3-6 biological replicates per condition. Error bars indicate %SD (Standard Deviation). For qPCR analysis, normalized gene expression levels were analyzed using one-way ANOVA. Error bars represent the standard error of the mean (SEM).

#### Immunoprecipitation and Western blots

For Immunoprecipitation (IP) from S2R+ cells, protein for each condition was prepared by lysing 1 well of a 6-well dish, 4 days after transfection. 2mg/ml was used per IP experiments performed with camelid antibodies GFP-Trap Magnetic Agarose (Cat# gtma-20, Chromotek); RFP-Trap Magnetic Agarose (Cat# rtma-20, Chromotek); Myc-Trap magnetic agarose (Cat# ytma-20; ChromoTek); as per the manufacturer’s protocol.

Western blots were performed detailed in our prior publication 1. Antibodies used include Chicken anti-GFP (Cat#ab13970, Abcam) 1:2000 in TBS-0.05% tween-20 (TBS-T), Rabbit-anti Upd2 [custom antibody AR4398 generated at YenZym Inc., to Upd2 peptide 318-334: RRPRRNSAERRHLAAIHC; antibody was verified by its recognition of GFP tagged Upd2 pull down in western blots] 1:2000 in TBS-T 0.05%; Rabbit-anti-GRASP (Cat# ab 30315, Abcam) 1:10000 in 1% BSA TBS-T 0.1%; Rabbit-anti RFP (Cat# 600-401-379, Rockland) 1:2500 in0.1% TBS-T. The secondary antibodies used were HRP conjugated Goat antibodies directed to appropriate species. The western blot was developed using Amersham ECL kit Start or Prime (Cat# RPN3243 or # RPN2232, GE life sciences).

#### Tissue preparation for fixed immunostaining

For fat body preparation, incisions using dissection scissors (Cat# 500086, World Precision Instruments Inc) were made to release the ventral abdomen from the rest of the fly body. Flies used for dissection were adult males, 7-10 days old. Dissections were done in Ringer’s medium (1.8 mM CaCl2, 2mM Kcl, 128mM NaCl, 4mM MgCl2. 35mM sucrose, 5mM HEPES). The tissue was fixed in 4% formaldehyde for 20 minutes and rinsed with PBS. For immunohistochemistry, the fixed fat tissue was permeabilized in PBS+ 1.0% Triton-X-100 for 3X washes 5 minutes, subsequently washed with PBS+0.3% Triton-X-100 (Fat wash). Blocked for 30 minutes at room temperature (RT) with gentle agitation in Fat wash+ 5% Normal donkey serum (Block). Primary antibodies, diluted in block, included Rabbit-anti-RFP (1:500, Rockland, #600-401-379); Chicken-anti-GFP (1:2000, Abcam, #ab13970); Mouse-anti-HA (1:200; Abcam, #ab18181); mouse-anti-Lamin (1:100; ADL67.10 DSHB); Rabbit-anti-GABARAP (Atg8) (1:500; Abcam, # ab109364); and incubated overnight at 4C.

Washed 3X-5X for 15 mins each the following day in the fat wash at (RT) incubated with appropriate secondary antibody (from Jackson Immunoresearch) at a concentration of 1:500 in the block for 2-4 hours at RT. Washed 3X-5X for 5-15 minutes in the fat wash, mounted in SlowFade Gold antifade reagent with DAPI (Cat# S36938, Invitrogen). Images were captured with Zeiss LSM 800 confocal system and analyzed with Zeiss ZenLite or ImageJ.

#### Cell staining

For immunohistochemistry, cells were fixed for 20 minutes in 4% formaldehyde, washed in PBS for 5 quick changes, permeabilized in PBS+ 0.1% Triton-X-100 for 3X washes 5 minutes, subsequently washed with PBS+0.1% Triton-X-100 (Cell wash). Blocked for 30 minutes at room temperature (RT) with gentle agitation in Cell wash+ 5% Normal donkey serum (Block). Primary antibodies – Rabbit-anti-RFP (1:500, Rockland, #600-401-379); Chicken-anti-GFP (1:2000, Abcam, #ab13970); mouse-anti-Lamin (1:100; ADL67.10 DSHB) – were diluted in block incubated overnight at 4oC. Washed 3X-5X for 15 mins each was the following day in the fat wash at (RT) incubated with appropriate secondary antibody at a concentration of 1:500 (from Jackson Immunoresearch) in the block for 2-4 hours at RT. Washed 3X-5X for 5-15 minutes in cell wash, mounted with SlowFade Gold antifade reagent with DAPI (Cat# S36938, Invitrogen). Images were captured with Zeiss LSM 800 or Zeiss Elyra 7 with Lattice SIM confocal systems and analyzed with Zeiss ZenLite or ImageJ 14.

#### Hunger-driven feeding

For hunger-driven feeding analysis, age-matched W1118 flies were given a normal diet or HSD for 5, 7, 14, 21, 24, and 28 days after an initial 7 days of development on a normal diet. All other experiments were performed for 7-day or 14-day durations. Sixteen hours before feeding behavior assessment, half of the flies from each treatment were moved to starvation media. Individual flies were placed in a single well of fly liquid-food interaction counter (FLIC) and supplied with a 1% sucrose liquid diet. Detailed methods for how FLIC operates can be found in Ro et al., 2014 [103]. Fly feeding was measured for the first three hours in the FLIC, and all FLICs were performed at 10 am local time. For each FLIC, half of the wells (n=6/FLIC) contained the fed group, and the other half contained the starved group of flies for direct comparison. 12-30 flies were measured for analysis of feeding and normalized to the control fed group as a percentage. A signal above 40 (a.u.) was considered a feeding event. Analysis of feeding events was performed using R.

#### Triglyceride Measurements

TAG assays were carried out as previously described in (Rajan et al., 2017). In brief: Flies were homogenized in PBST (PBS + 0.1% Triton X-100) using 1mm zirconium beads (Cat#ZROB10, Next Advance) in a Bullet Blender® Tissue homogenizer (Model BBX24, Next Advance). Samples were heated to 70°C for 10 minutes, then centrifuged at 14,000 rpm (in a refrigerated tabletop centrifuge). 10.0 μl of the supernatant was applied to determine the level of TAG in the sample, using the following reagents obtained from Sigma: Free glycerol (cat # F6428-40ML), Triglyceride reagent (cat# T2449-10ML), and Glycerol standard (cat# G7793-5ML). Three adult males were employed per biological replicate. Note: For adult TAG assays, the most consistent results, with the lowest standard deviations, were obtained with 10-day old males. TAG readings from whole fly lysate (n=4 replicates of 3 flies each) were normalized to the number of flies per experiment. The normalized ratio from the control served as a baseline, and data are represented as fold change of experimental genotypes with respect to the control.

#### Survival Assays

Survival curves were done using flies harvested in a 24-hour time frame and aged for 7 days. Ten males per vial were flipped onto 1% sucrose agar starvation food or a 30% High-sugar diet (HSD). The number of dead flies was recorded each day until every fly had died. Flies were kept in a 25°C incubator with a 12-hour light-dark cycle for the entirety of the experiment. Survival analysis was performed using the Survival Curve module of GraphPad Prism. A Mantel-cox test was the test used to determine statistical significance. Greater than 90 flies were used per genotype per curve. Data consolidated from 3 independent experiments is shown.

#### Image acquisition, Morphometrics and image analysis

All cell biological experiments in fat cells and S2R+ cells, were perform 3 independent times at least. For fat imaging 6-8 fat explants per experimental condition was used. For cells, at least 10 Z-stacks, with 5-10 cells per field of view was captured per experimental condition. The confocal settings for laser power and gain were maintained consistent per experiment across different genotypes or dietary conditions. To prevent bias, the person performing the starved versus fed imaging was blinded to the sample’s status. Excel or GraphPad Prism 7 software was used for data quantification and generation of graphs. Quantification of nuclear localization in S2R+ was performed as follows:

Image analysis pipelines were built in MATLAB R2020b, and the associated scripts are available on request to the lead author. Three types of analysis were performed in 3D to quantify 1- the nucleus-to-whole cell intensity ratio of Upd2 in S2 cells, 2- the vesicle-to-whole cell intensity of Atg8 in S2 cells, and 3- the mean nuclear pixel intensity of Upd2 or Atg8 in fat cells.

To generate the 3D nuclear masks, Lamin or DAPI stacks were first maximally projected along the z-axis. After global thresholding, basic morphological operations, and watershed transform, the locations of the nuclear centroids in the x-y plane were used to scan the z-stacks and reconstruct the nuclear volume. The whole-cell volume was approximated from the nuclear volume by dilation. Atg8 vesicles were segmented within each cell volume by performing morphological filtering and image opening with defined structural elements. The accuracy of the segmentation was assessed by manual inspection of random cells.

Once these cellular compartments had been defined, they were used as masks to extract intensity values of the signal of interest and to compute either the mean pixel intensity in the nucleus, the ratio of the integral intensities between the nucleus and the whole cell, or between the combined vesicles and the whole cell. Two-sided Wilcoxon rank-sum tests were used to assess the statistical significance of pairwise comparisons between experimental conditions.

We determined that our data-points were normally distributed, based on two measures: i) A GraphPad outlier test did not identify any outliers in our data; and ii) the majority of our data points for a particular condition were relatively similar to one other, with only a small standard error of mean or standard deviation.

#### RNAseq

*RNA prep:* Total RNA prepared from 30 flies per genotype in triplicates, using the Direct-zol RNA miniprep kit (Zymo Research, cat#R2071). cDNA prepared with iScript cDNA Synthesis (Bio-Rad, cat#1708891), and 1mg RNA applied per reaction.

*RNA quality control:* Total RNA integrity was checked using an Agilent 4200 TapeStation (Agilent Technologies, Inc., Santa Clara, CA) and quantified using a Trinean DropSense96 spectrophotometer (Caliper Life Sciences, Hopkinton, MA).

*RNA-seq Expression Analysis:* RNA-seq libraries were prepared from total RNA using the TruSeq Stranded mRNA kit (Illumina, Inc., San Diego, CA, USA). Library size distribution was validated using an Agilent 4200 TapeStation (Agilent Technologies, Santa Clara, CA, USA). Additional library QC, blending of pooled indexed libraries, and cluster optimization were performed using Life Technologies’ Invitrogen Qubit® 2.0 Fluorometer (Life Technologies-Invitrogen, Carlsbad, CA, USA). RNA-seq libraries were pooled (35-plex) and clustered onto an SP flow cell. Sequencing was performed using an Illumina NovaSeq 6000 employing a paired-end, 50 base read length (PE50) sequencing strategy.

*RNA-seq analysis:* STAR v2.7.1[104] with 2-pass mapping was used to align 50bp paired-end reads to Drosophila melanogaster genome build BDGP6.32 and Ensembl gene annotation 104 (http://uswest.ensembl.org/Drosophila_melanogaster/Info/Index). FastQC 0.11.9 (https://www.bioinformatics.babraham.ac.uk/projects/fastqc/) and RSeQC 4.0.0 [105] were used for QC, including insert fragment size, read quality, read duplication rates, gene body coverage, and read distribution in different genomic regions. FeatureCounts [106] in Subread 1.6.5 was used to quantify gene-level expression by strand-specific paired-end read counting. Bioconductor package edgeR 3.26.8 (https://academic.oup.com/bioinformatics/article/26/1/139/182458) was used to detect differential gene expression between conditions. Genes with low expression were excluded by requiring at least one count per million in at least N samples (N is equal to the number of samples in the smallest group). The filtered expression matrix was normalized by the TMM method [107] and subject to significance testing using a quasi-likelihood pipeline implemented in edgeR. Genes were deemed differentially expressed if absolute fold changes were above 2 and Benjamini-Hochberg adjusted p-values were less than 0.01.

---

**Figure S1:**
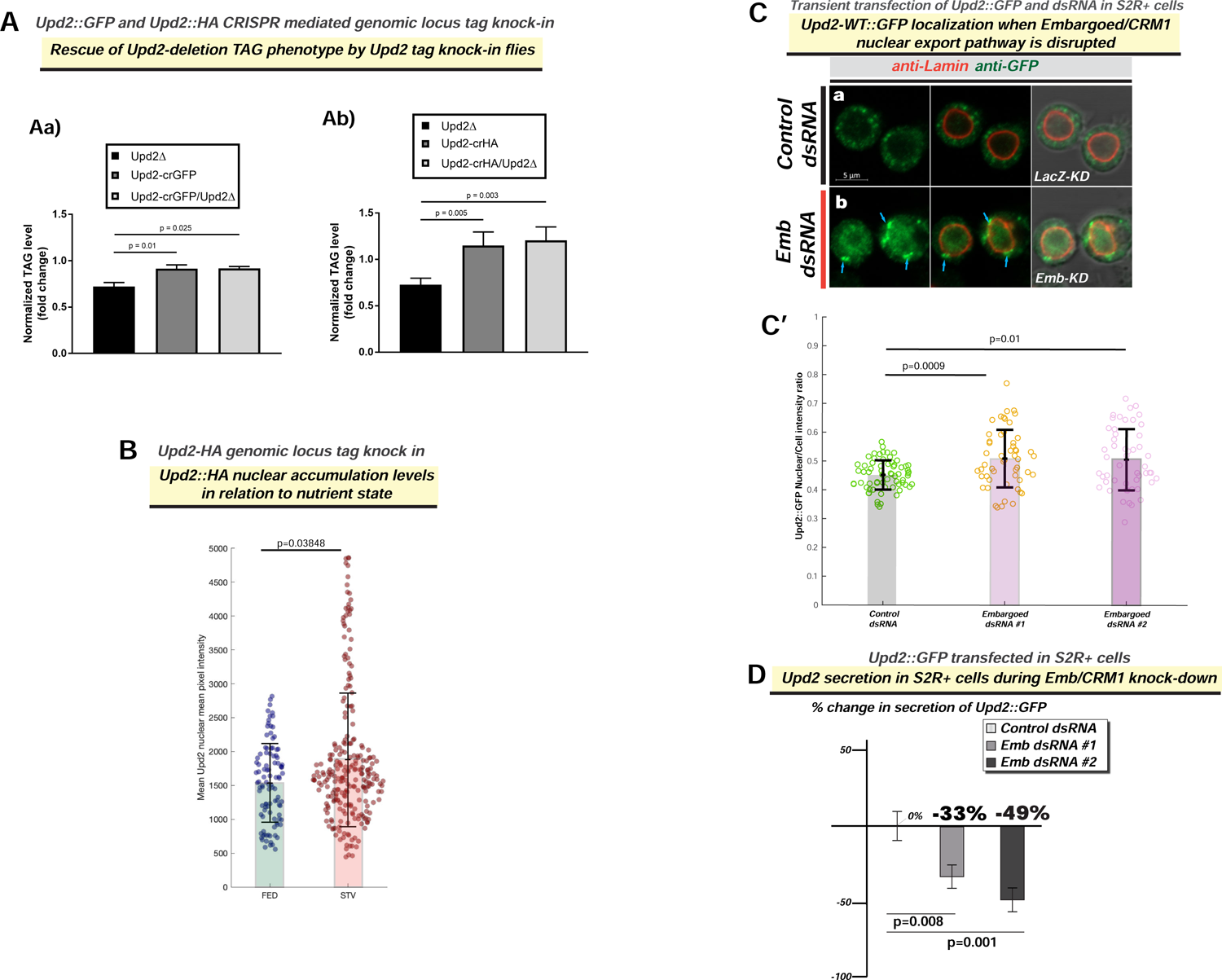
Companion to Main Figure 1. (A) Rescue of upd2Δ triglyceride (TAG) storage reduction phenotype by endogenous CRISPR tag knock-in of HA [Upd2-HA] (S1B) and GFP [Upd2-GFP] (1A) in the Upd2 locus. (B) Upd2-HA endogenous nuclear accumulation is quantified using 3D-volumetric segmentation-based calculations (See Methods). Compare with similar results observed for a GFP-tagged Upd2 in Main Figure 1A’. (C) Confocal micrographs of single optical-slices of *Drosophila* S2R+ cells transiently co-transfected with Upd2-WT::GFP (green) stained with Lamin (red). In fed state, cells are co-transfected with (a) LacZ-dsRNA (control), or (b) CRM1/Embragoed (Emb)-KD (Emb dsRNA). Emb-KD increases Upd2-WT::GFP nuclear accumulation and shows an enrichment of Upd2-WT::GFP at the interface of nuclear lamina and cytoplasm (see blue arrows). Scale bar is 5µm. Two independent *Emb-dsRNAs* (#1 and #2) were used. For B, C’ statistical significance is quantified by the two-sided Wilcoxon test. Each dot represents a fat cell nucleus (B) or cell ( C’). 50-100 replicates were quantified per genotype per condition. (D) Normalized percent fold change in secreted GFP signal detected by GFP sandwich ELISA assay performed on conditioned media of S2R+ cells transiently transfected with Upd2-WT::GFP dsRNA for control (LacZ) or two independent *Emb dsRNAs*. ELISA for Upd2-WT::GFP was performed after 5 days of knockdown, and the amount of Upd2-WT::GFP in a 24-hour period (Day 5) was assessed. Statistical significance is quantified by unpaired two-tailed t-test on 6 biological replicates per condition.

**Figure S2:**
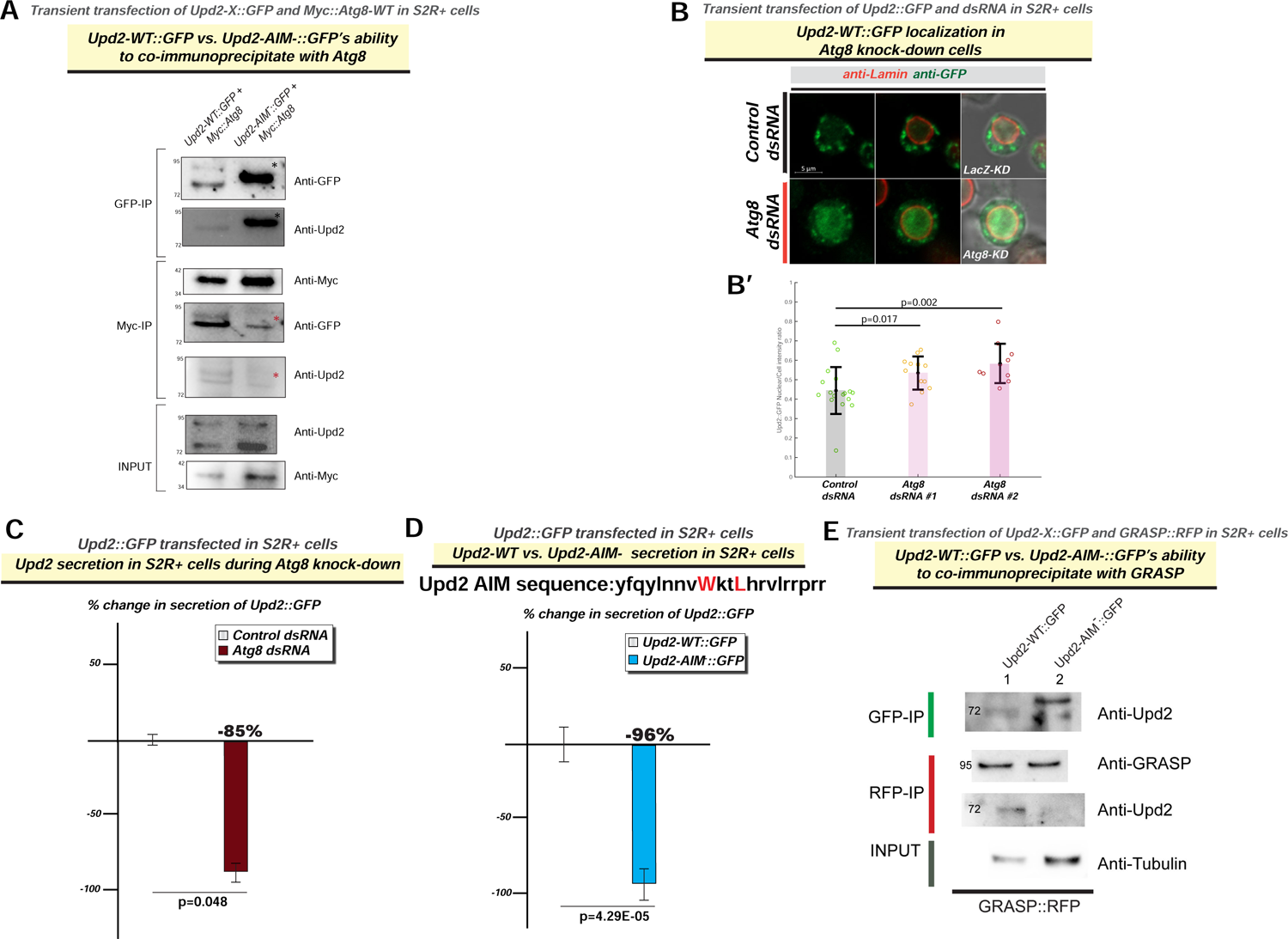
Companion to Main Figure 2. (A) GFP- and Myc-IPs were prepared from S2R+ cells transiently transfected with the indicated cDNAs. GFP, Myc IPs and 2% of the input were analyzed by immunoblotting for the indicated proteins. Note Upd2-AIM-always runs slightly higher than Upd2 (see asterisk) and is always more abundant in the lysate as the Upd2-AIM-is not secreted (see D below), but Upd2-AIM-it is not readily detectable in the Myc-IPs (see red asterisk), despite being more abundant in input. Atg8-Myc is always more abundant in Upd2-AIM-input than in Upd2-WT input, despite loading same amount of protein in input. (B) Confocal micrographs of single optical-slices of *Drosophila* S2R+ cells transiently co-transfected with Upd2-WT::GFP (green) stained with Lamin (red). (a) In fed state, in cells co-transfected with LacZ-dsRNA (control) Upd2-WT::GFP is localized to cytosolic puncta and nuclear localization is not obvious. (b) In fed state, Atg8-KD (Atg8 dsRNA) significant increase in Upd2-WT::GFP nuclear accumulation is observed. Scale bar is 5um and right most panel shows DIC image merge. In B’, the ratio of Upd2::GFP nuclear/whole cell intensity is plotted. Each dot represents a cell, 20-40 cells were counted per dsRNA condition. Two independent Atg8-dsRNAs were used. Statistical significance is quantified by two-sided Wilcoxon sum rank tests. (C) Normalized percent fold change in secreted GFP signal detected by GFP sandwich ELISA assay performed on conditioned media of *Drosophila* S2R+ cells transiently transfected with Upd2-WT::GFP dsRNA for control (LacZ) or Atg8 dsRNA. ELISA for Upd2-WT::GFP was performed after 5 days of knockdown, and the amount of Upd2-WT::GFP in a 24-hour period (Day 5) was assessed. Statistical significance is quantified by unpaired two-tailed t-test on 6 biological replicates per condition. (D) Normalized percent fold change in secreted GFP signal detected by GFP sandwich ELISA assay performed on conditioned media of S2R+ cells transiently transfected with Upd2-WT::GFP or Upd2-AIM-::GFP. Mutated residues in Upd2’s AIM is shown in red. Statistical significance is quantified by unpaired two-tailed t-test on 6 biological replicates per condition. (E) GFP- and RFP-IPs were prepared from S2R+ cells transiently co-transfected with GRASP-RFP and Upd2-WT::GFP or Upd2-AIM-::GFP. GFP, RFP IPs and 2% of the input were analyzed by immunoblotting for the indicated proteins. Note Upd2-AIM-always runs slightly higher than Upd2 and is always more abundant in the lysate as the Upd2-AIM- is not secreted, but it is not readily detectable in the RFP-IPs, despite loading more input.

**Supplemental Figure 3:**
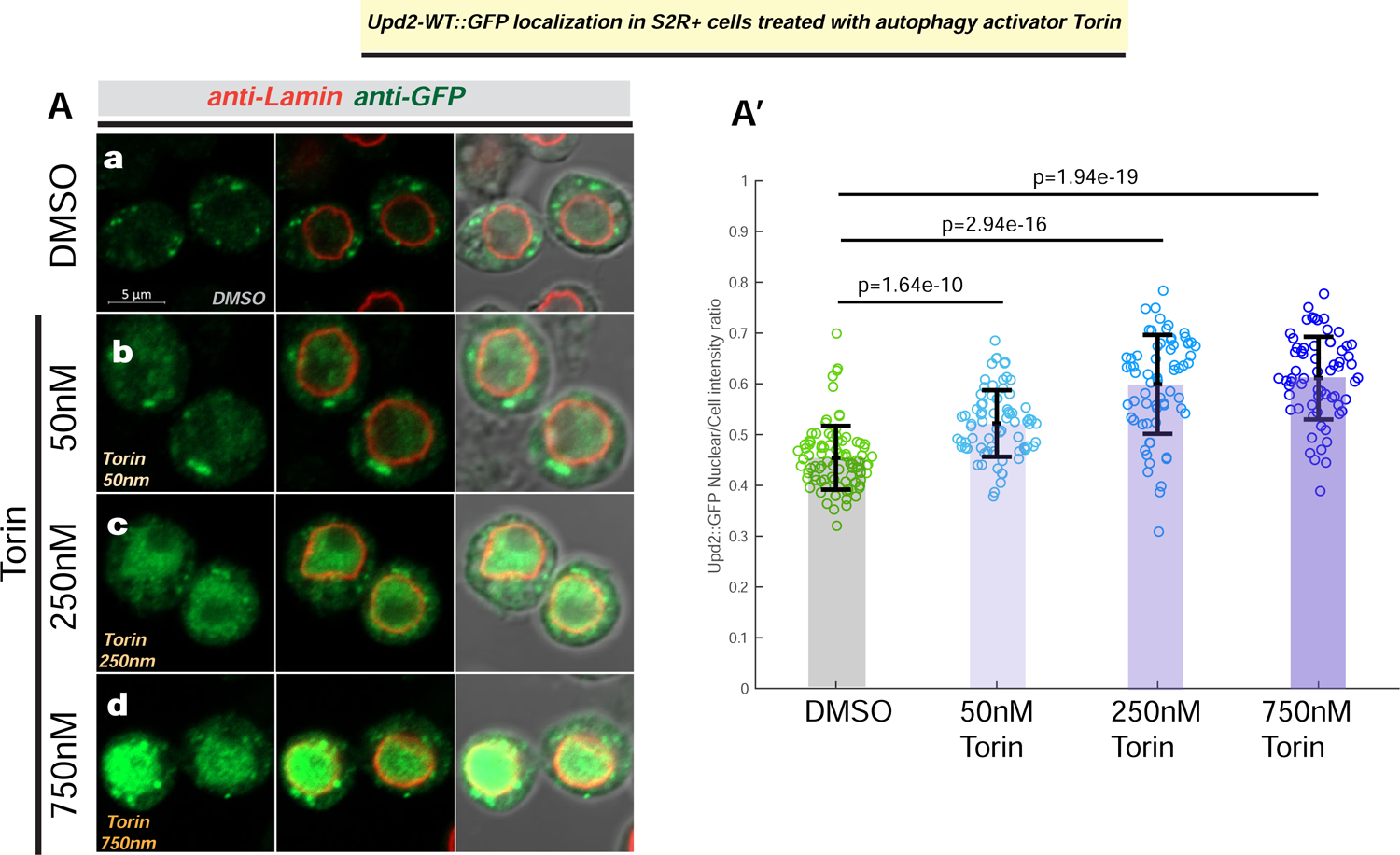
Companion to Main Figure 3. (A) Confocal micrographs of single optical-slices of *Drosophila* S2R+ cells transiently transfected with Upd2-WT::GFP (green) stained with Lamin (red) and treated with DMSO (a) or indicated concentrations of autophagy activator Torin1 (b, c, d). Scale bar is 5µm and right most panel shows DIC image merge. In A’, the ratio of Upd2::GFP nuclear/whole cell intensity is plotted. Each dot represents a cell, 50-100 cells were counted per condition. Statistical significance is quantified by unpaired two-tailed t-test on 6 biological replicates per condition.

**Figure S4:**
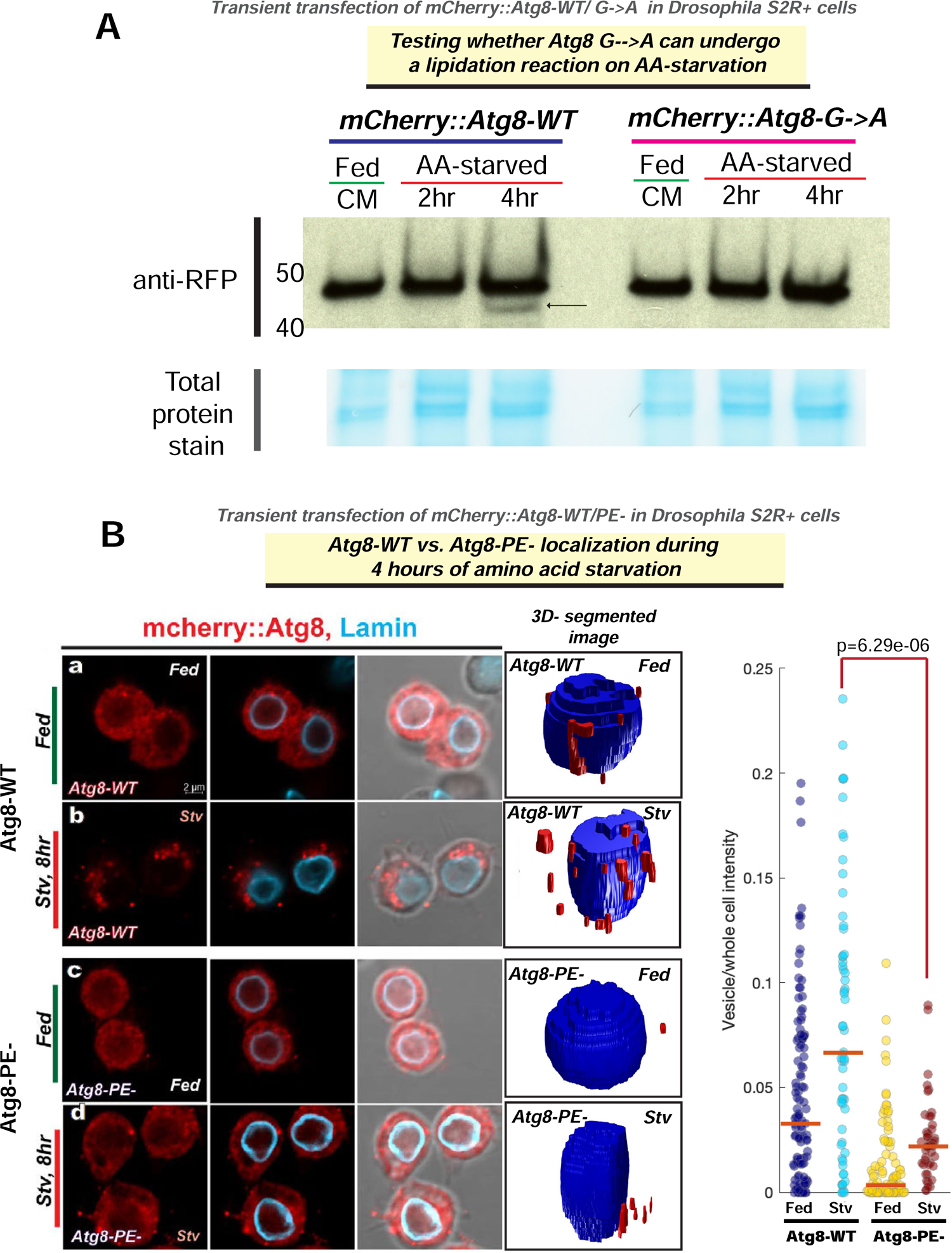
Companion to Main Figure 3. (A) Western blot on lysates prepared from cells transfected with mCherry::Atg8-WT or mCherry::Atg8-G◊ A. Cells were either maintained in complete media (Fed-CM) or AA-acid starved for 2 hours (2hr) and 4 hours (4hr). The cell lysates were probed with anti-RFP antibody. Atg8 (15kDa Atg8+ 27kDa mCherry) is recognized at ∼42kDa. On 4 hr AA starvation, only in Atg8-WT, a lower doublet band is observed (see arrow), but not in the mCherry::Atg8 G◊A, suggesting defective lipidation. Loading is shown as a total protein stain for the blot. (B) Confocal micrographs of single optical-slices of *Drosophila* S2R+ cells transiently transfected with (a, b) mCherry::Atg8-WT (red) or (c, d) mCherry::Atg8-PE-(red) stained with Lamin (blue) in either well-fed state (a, c) or AA-starved (b, d). Scale bar is 2um and right most panel shows DIC image merge. Atg8 vesicles were segmented within each cell volume by performing morphological filtering (See Methods). Representative images of thresholding analyses corresponding to confocal images are shown in right most panel. Red represents the vesicular segmentation. Portion of Atg8 signal trapped in the vesicles, defined as the ratio between Atg8 vesicle intensity and whole cell is quantified. Each dot represents one cell. P-value calculated using the two-sided Wilcoxon rank sum tests. Note the significant increase in Atg8-WT vesicular/whole cell ratio during starvation is not observed in Atg8-PE-. Atg8-PE-vesicle amounts phenocopy Atg8-WT in the fed state.

**Figure S5:**
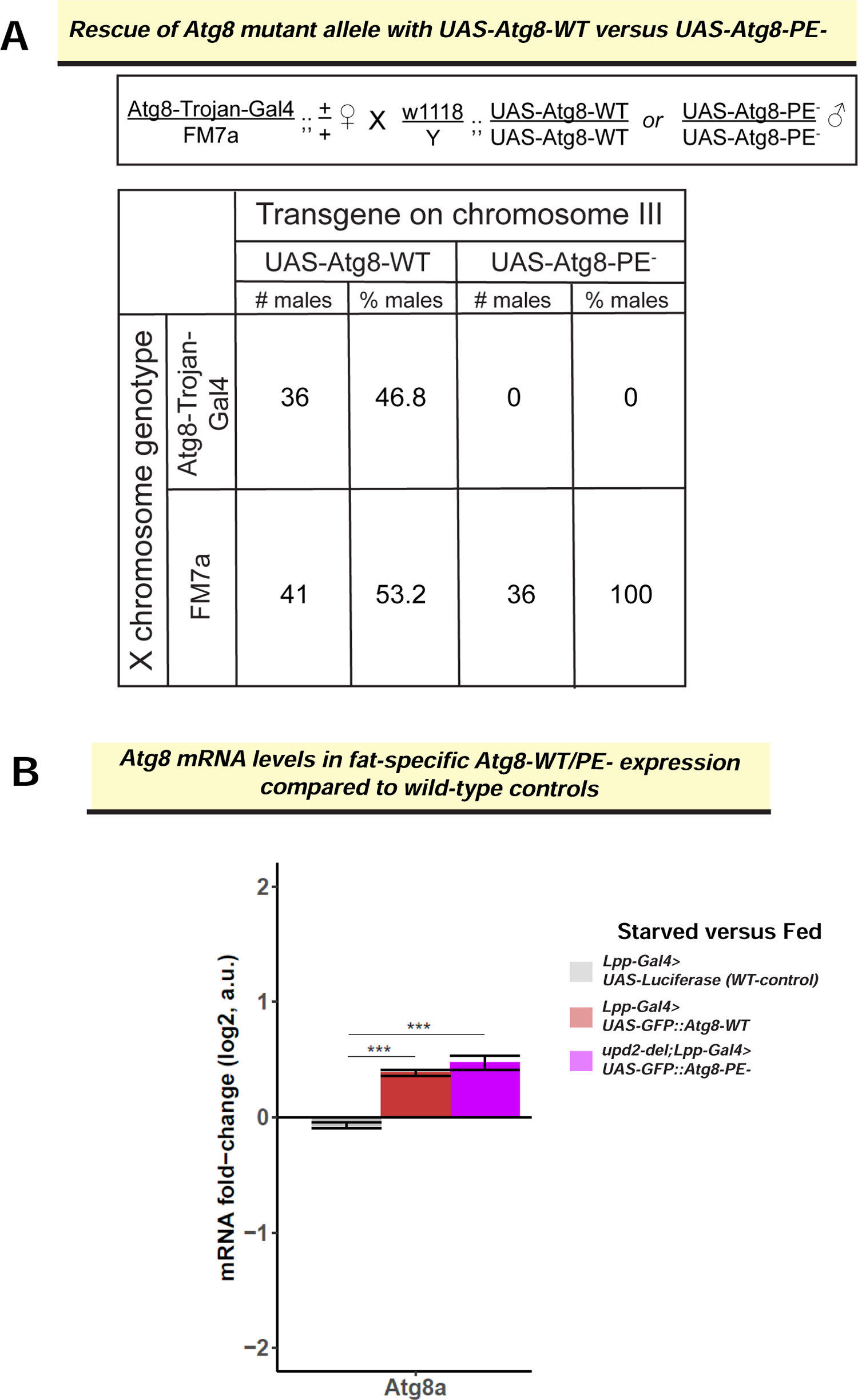
Companion to Main Figure 4. (C) Assessing the rescue of an Atg8 knock out allele-*Atg8-Trojan-Gal4* (see methods), by the UAS-Atg8 transgenes. In the text box is outlined the crossing scheme depicting the Atg8-Trojan-Gal4 allele rescue with wild-type Atg8 transgene (UAS-GFP::Atg8-WT) or the lipidation-defective Atg8 transgene (UAS-GFP::Atg8-PE-). Table shows the number of progenies counted from the cross and the % of transgenic rescue. While the expected percentage of flies with UAS-GFP-Atg8-WT, as expected the UAS-Atg8-PE-(lipidation-defective version) does not rescue the Atg8 knockout allele. (D) Atg8 mRNA levels assessed by RNAseq in the indicated genotypes. Each bar represents difference in fold change between starved and fed state of the indicated genotype. Error bars represent the standard deviation of log2 fold change among biological replicates (n=3). Significance was calculated using t-test and followed by *post-hoc* analyses of q-values to account for false discovery rate (See Methods). Values of q<0.05 were considered significant. Atg8 transgenes (red and violet bars) are upregulated on starvation and expressed at higher levels than control (grey). The difference between Atg8-WT and Atg8-PE-expression is not significant.

**Figure S6:**
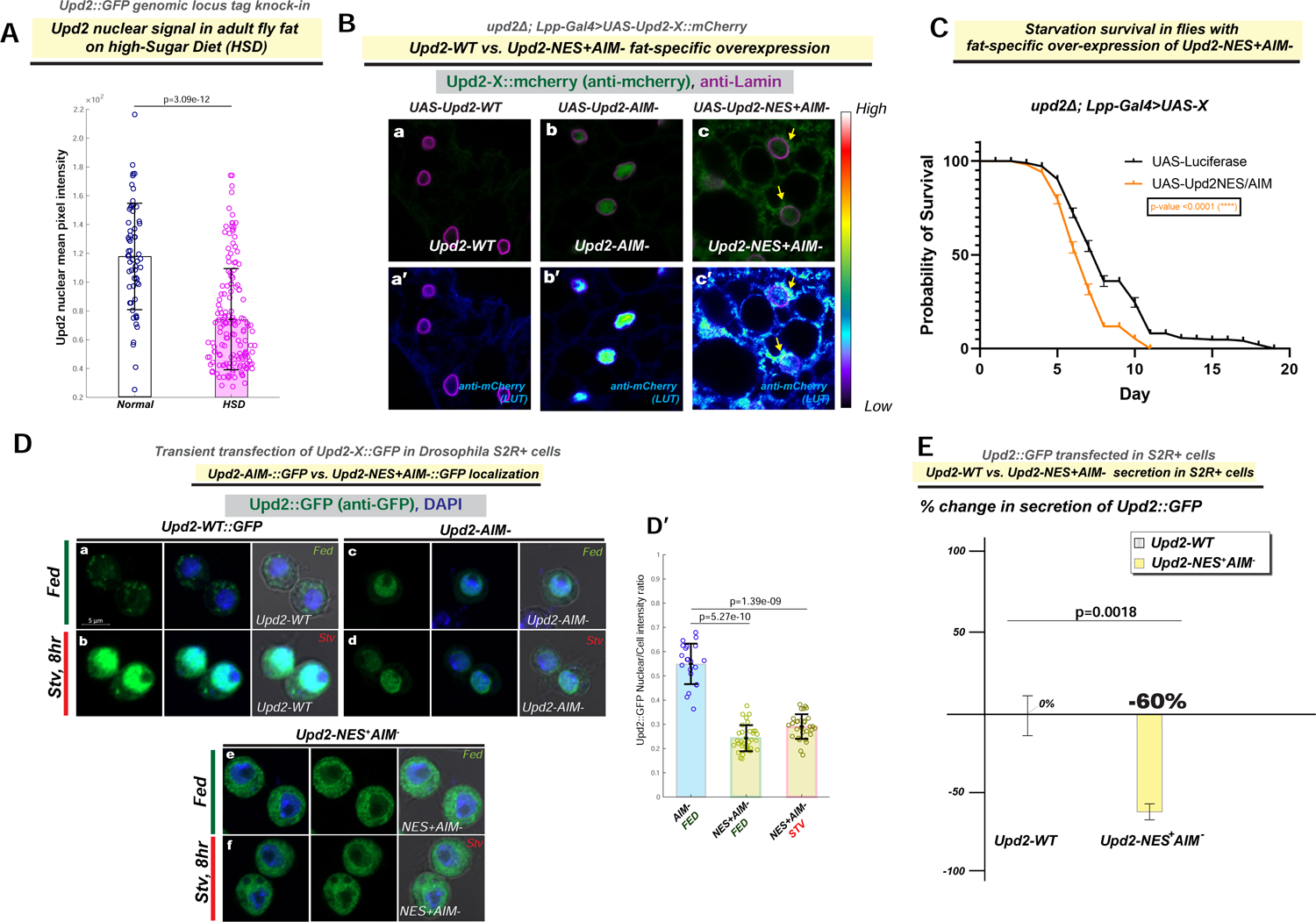
Companion to Main Figure 5. (A) Confocal micrographs of single optical sections of fixed and immunostained adult fly fat from transgenic flies expressing mCherry tagged Upd2-cDNAs for (a, a’) Upd2-WT (*UAS-Upd2-WT*); (b, b’) Upd2’s AIM mutated (*UAS-Upd2-AIM-*); (c,c’) Upd2-AIM-engineered with a N-terminal canonical nuclear export signal (NES) (*UAS-Upd2-NES+-AIM-*) under the control of a fat-cell specific promoter (Lpp-Gal4) in a upd2-deletion background (*upd2Δ; Lpp-Gal4*). In Upd2-WT in fed state is detectable in very low levels in the nucleus (see look up table (LUT)) and yellow arrows in a Upd2-AIM-shows significantly nuclear accumulation even in a fed state (b’). Whereas addition of NES+ to Upd2-AIM (c,c’) causes Upd2 to be localized in a diffuse pattern in the cytosol and localize to periphery of nuclear lamina see arrows. (B) Flies expressing cDNAs (UAS-X) for *Upd2-NES+AIM-* and control (*UAS-Luciferase*) specifically in fly fat in upd2 deletion background (*upd2Δ Lpp-Gal4>UAS-X*). Transgenic flies with fat-specific overexpression of AIM- that has a canonical NES (Upd2-NES+-AIM-) results in starvation-sensitivity in comparison to controls as assessed by the Log-rank (Mantel-Cox) test. See also Main Figure 5B on characterization of starvation sensitivity of *UAS-Upd2-AIM-*. (C) Confocal micrographs of single optical-slices of *Drosophila* S2R+ cells transiently co-transfected with Upd2::GFP variants (a, b) WT; (c, d) (e, f) NES+AIM-Upd2::GFP (green) stained with DAPI (blue) and anti-GFP (green) in a fed state (a,c,) or AA-starved for 8 hours (b, d). Scale bars are 5um right most panel shows DIC image merge. Note that addition of NES+ renders Upd2-AIM-cytosolic in both fed and starved states. Note also that unlike Upd2::WT which are localized to punctate in fed states(Ca), the addition of NES+ to Upd2-AIM-display a diffuse cytosolic localization (Ce). C’ Quantification of nuclear to cytosolic ratio of Upd2-GFP based on 3D-volumetric methods, to assess the difference between Upd2-AIM- in fed state versus Upd2-NES+-AIM-. Upd2-NES+AIM- is significantly less nuclear than Upd2-AIM- in both fed and starved states. (D) Normalized percent fold change in secreted GFP signal detected by GFP sandwich ELISA assay performed on conditioned media of S2R+ cells transiently transfected with Upd2::GFP variants: WT; NES+AIM- the amount of Upd2-WT::GFP in a 24-hour period was assessed. Statistical significance is quantified by unpaired two-tailed t-test on 6 biological replicates per condition. Note that despite being localized to the cytosol Upd2-NES+-AIM- is not secreted, consistent with the requirement for Atg8 during secretion.

**Figure S7:**
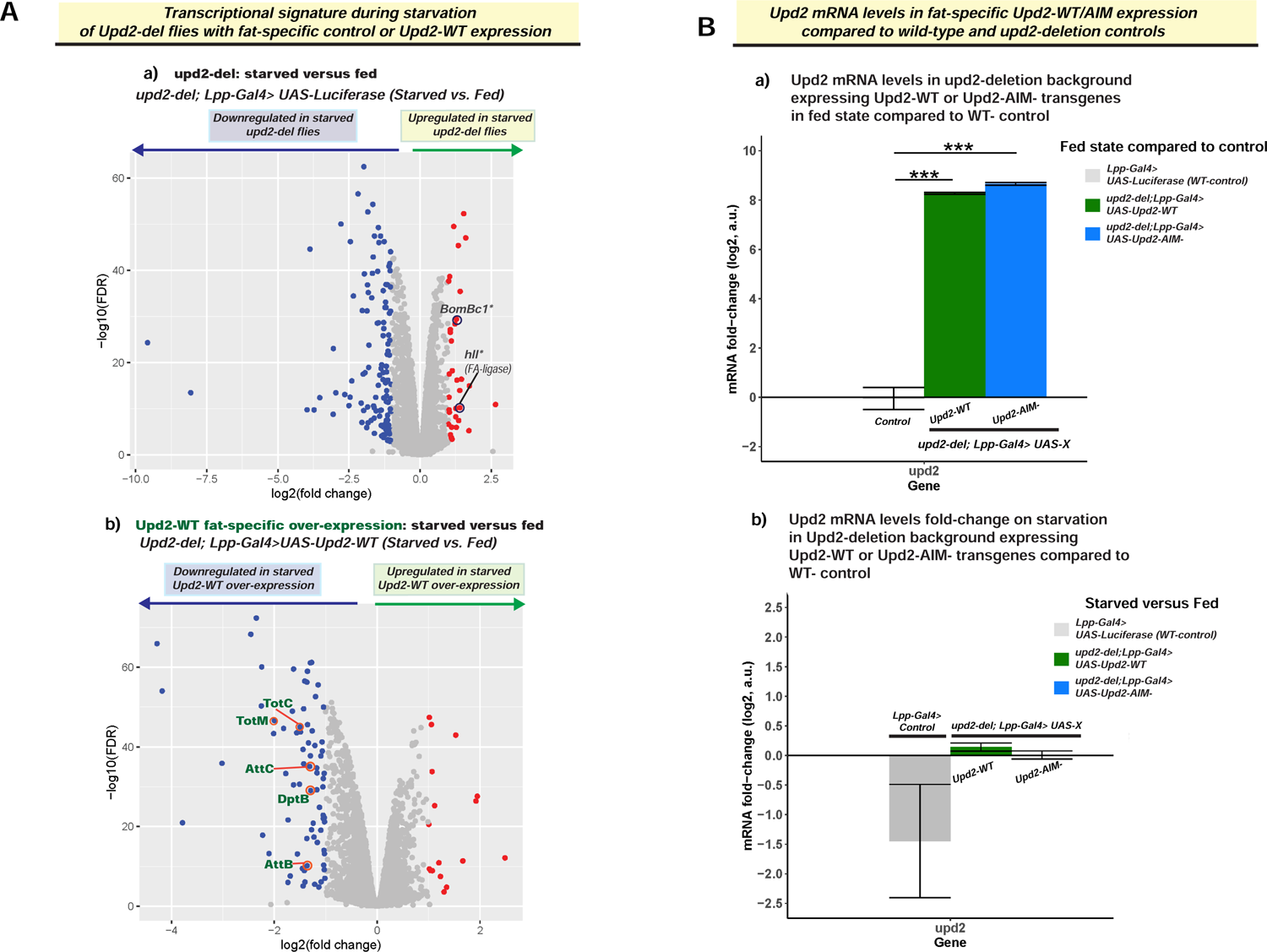
Companion to Main Figure 6. (A) Volcano plot depicts starvation-induced changes in transcriptome, as assayed by RNA-seq (see methods), of control (a; *upd2Δ-Lpp-Gal4> UAS-Luciferase*) and Upd2-WT-fat specific overexpression (b; *upd2Δ; Lpp-Gal4> UAS-WT-*). Genes that are significantly upregulated during starvation are shown as red dots and downregulated genes as blue dots. In the control flies (a), among the six genes upregulated on starvation, 2 genes encode genes involved in innate immunity IBIN and SPH93 are not upregulated (See Main Figure 6B). (B) Upd2 mRNA levels assessed by RNAseq in the indicated genotypes. (a) Each bar represents difference in fold change between fed state control (Lpp-Gal4> UAS-Luciferase) and either Upd2-WT (green) or Upd2-AIM-overexpression in the WT background. Error bars represent the standard deviation of log2 fold change among biological replicates (n=3). Significance was calculated using t-test and p<0.05 were considered significant. *** represents p<0.0001. Note Upd2-AIM- and Upd2-WT expression levels are comparable. (b) Each bar represents difference in fold change between starved and fed state of the indicated genotype. Error bars represent the standard deviation of log2 fold change among biological replicates (n=3).

## References

1. Leopold, P. and N. Perrimon, Drosophila and the genetics of the internal milieu. Nature, 2007. 450(7167): p. 186-8.

2. Efeyan, A., W.C. Comb, and D.M. Sabatini, Nutrient-sensing mechanisms and pathways. Nature, 2015. 517(7534): p. 302-10.

3. Singh, R. and A.M. Cuervo, Autophagy in the cellular energetic balance. Cell Metab, 2011. 13(5): p. 495–504.

4. Flier, J.S., Starvation in the Midst of Plenty: Reflections on the History and Biology of Insulin and Leptin. Endocr Rev, 2019. 40(1): p. 1–16.

5. Neufeld, T.P. and E.H. Baehrecke, Eating on the fly: function and regulation of autophagy during cell growth, survival and death in Drosophila. Autophagy, 2008. 4(5): p. 557–62.

6. Zirin, J. and N. Perrimon, Drosophila as a model system to study autophagy. Semin Immunopathol, 2010. 32(4): p. 363–72.

7. Rajan, A. and N. Perrimon, Of flies and men: insights on organismal metabolism from fruit flies. BMC Biol, 2013. 11: p. 38.

8. Droujinine, I.A. and N. Perrimon, Interorgan Communication Pathways in Physiology: Focus on Drosophila. Annu Rev Genet, 2016. 50: p. 539–570.

9. Kim, S.K., et al., Discovering signaling mechanisms governing metabolism and metabolic diseases with Drosophila. Cell Metab, 2021. 33(7): p. 1279–1292.

10. Baker, K.D. and C.S. Thummel, Diabetic larvae and obese flies-emerging studies of metabolism in Drosophila. Cell Metab, 2007. 6(4): p. 257–66.

11. King-Jones, K. and C.S. Thummel, Nuclear receptors--a perspective from Drosophila. Nat Rev Genet, 2005. 6(4): p. 311–23.

12. Musselman, L.P. and R.P. Kuhnlein, Drosophila as a model to study obesity and metabolic disease. J Exp Biol, 2018. 221(Pt Suppl 1).

13. Teleman, A.A., I. Ratzenbock, and S. Oldham, Drosophila: a model for understanding obesity and diabetic complications. Exp Clin Endocrinol Diabetes, 2012. 120(4): p. 184–5.

14. Tennessen, J.M., et al., Methods for studying metabolism in Drosophila. Methods, 2014. 68(1): p. 105–15.

15. Gillette, C.M., J.M. Tennessen, and T. Reis, Balancing energy expenditure and storage with growth and biosynthesis during Drosophila development. Dev Biol, 2021. 475: p. 234–244.

16. van Dam, E., et al., Sugar-Induced Obesity and Insulin Resistance Are Uncoupled from Shortened Survival in Drosophila. Cell Metab, 2020. 31(4): p. 710–725 e7.

17. Ahima, R.S., et al., role of leptin in the neuroendocrine response to fasting. Nature, 1996. 382(6588): p. 250-2.

18. Farooqi, I.S. and S. O’Rahilly, Leptin: a pivotal regulator of human energy homeostasis. Am J Clin Nutr, 2009. 89(3): p. 980S–984S.

19. Rajan, A. and N. Perrimon, Drosophila Cytokine Unpaired 2 Regulates Physiological Homeostasis by Remotely Controlling Insulin Secretion. Cell, 2012. 151(1): p. 123–137.

20. Kolaczynski, J.W., et al., Responses of leptin to short-term fasting and refeeding in humans: a link with ketogenesis but not ketones themselves. Diabetes, 1996. 45(11): p. 1511–5.

21. Kolaczynski, J.W., et al., response of leptin to short-term and prolonged overfeeding in humans. J Clin Endocrinol Metab, 1996. 81(11): p. 4162–5.

22. Hickey, M.S., et al., leptin is related to body fat content in male distance runners. Am J Physiol, 1996. 271(5 Pt 1): p. E938-40.

23. Brent, A.E. and A. Rajan, Insulin and Leptin/Upd2 Exert Opposing Influences on Synapse Number in Fat-Sensing Neurons. Cell Metab, 2020. 32(5): p. 786–800 e7.

24. Friedman, J.M. and J.L. Halaas, Leptin and the regulation of body weight in mammals. Nature, 1998. 395(6704): p. 763-70.

25. Ertekin, D., et al., Down-regulation of a cytokine secreted from peripheral fat bodies improves visual attention while reducing sleep in Drosophila. PLoS Biol, 2020. 18(8): p. e3000548.

26. Flier, J.S. and E. Maratos-Flier, Leptin’s Physiologic Role: Does the Emperor of Energy Balance Have No Clothes? Cell Metab, 2017. 26(1): p. 24–26.

27. Rajan, A., et al., A Mechanism Coupling Systemic Energy Sensing to Adipokine Secretion. Dev Cell, 2017. 43(1): p. 83–98 e6.

28. Kaushik, S., R. Singh, and A.M. Cuervo, Autophagic pathways and metabolic stress. Diabetes Obes Metab, 2010. 12 Suppl 2: p. 4–14.

29. Singh, R., et al., autophagy regulates lipid metabolism. Nature, 2009. 458(7242): p. 1131-5.

30. He, C. and D.J. Klionsky, Regulation mechanisms and signaling pathways of autophagy. Annu Rev Genet, 2009. 43: p. 67–93.

31. Singh, R., et al., autophagy regulates adipose mass and differentiation in mice. J Clin Invest, 2009. 119(11): p. 3329–39.

32. Shpilka, T., et al., Atg8: an autophagy-related ubiquitin-like protein family. Genome Biol, 2011. 12(7): p. 226.

33. Kirisako, T., et al., Formation process of autophagosome is traced with Apg8/Aut7p in yeast. J Cell Biol, 1999. 147(2): p. 435–46.

34. Huang, R., et al., Deacetylation of nuclear LC3 drives autophagy initiation under starvation. Mol Cell, 2015. 57(3): p. 456–66.

35. Ichimura, Y., et al., A ubiquitin-like system mediates protein lipidation. Nature, 2000. 408(6811): p. 488-92.

36. Nakatogawa, H., Y. Ichimura, and Y. Ohsumi, Atg8, a ubiquitin-like protein required for autophagosome formation, mediates membrane tethering and hemifusion. Cell, 2007. 130(1): p. 165–78.

37. Galluzzi, L. and D.R. Green, Autophagy-Independent Functions of the Autophagy Machinery. Cell, 2019. 177(7): p. 1682–1699.

38. Cunha, L.D., et al., LC3-Associated Phagocytosis in Myeloid Cells Promotes Tumor Immune Tolerance. Cell, 2018. 175(2): p. 429–441 e16.

39. Reggiori, F., et al., Coronaviruses Hijack the LC3-I-positive EDEMosomes, ER-derived vesicles exporting short-lived ERAD regulators, for replication. Cell Host Microbe, 2010. 7(6): p. 500–8.

40. Leidal, A.M., et al., The LC3-conjugation machinery specifies the loading of RNA-binding proteins into extracellular vesicles. Nat Cell Biol, 2020. 22(2): p. 187–199.

41. Heckmann, B.L., et al., LC3-Associated Endocytosis Facilitates beta-Amyloid Clearance and Mitigates Neurodegeneration in Murine Alzheimer’s Disease. Cell, 2019. 178(3): p. 536–551 e14.

42. Schneider, I., Cell lines derived from late embryonic stages of Drosophila melanogaster. J Embryol Exp Morphol, 1972. 27(2): p. 353–65.

43. Zacharogianni, M., et al., ERK7 is a negative regulator of protein secretion in response to amino-acid starvation by modulating Sec16 membrane association. EMBO J, 2011. 30(18): p. 3684–700.

44. Zacharogianni, M., et al., A stress assembly that confers cell viability by preserving ERES components during amino-acid starvation. Elife, 2014. 3.

45. Hombria, J.C., et al., Characterisation of Upd2, a Drosophila JAK/STAT pathway ligand. Dev Biol, 2005. 288(2): p. 420–33.

46. Wright, V.M., et al., Differential activities of the Drosophila JAK/STAT pathway ligands Upd, Upd2 and Upd3. Cell Signal, 2011. 23(5): p. 920-7.

47. Bard, F., et al., Functional genomics reveals genes involved in protein secretion and Golgi organization. Nature, 2006. 439(7076): p. 604-7.

48. Wendler, F., et al., A genome-wide RNA interference screen identifies two novel components of the metazoan secretory pathway. EMBO J, 2010. 29(2): p. 304–14.

49. Fasken, M.B., et al., A leptomycin B-sensitive homologue of human CRM1 promotes nuclear export of nuclear export sequence-containing proteins in Drosophila cells. J Biol Chem, 2000. 275(3): p. 1878–86.

50. Noda, N.N., Y. Ohsumi, and F. Inagaki, Atg8-family interacting motif crucial for selective autophagy. FEBS Lett, 2010. 584(7): p. 1379–85.

51. Zhang, M., et al., translocation of interleukin-1beta into a vesicle intermediate in autophagy-mediated secretion. Elife, 2015. 4.

52. Dupont, N., et al., Autophagy-based unconventional secretory pathway for extracellular delivery of IL-1beta. Embo J, 2011. 30(23): p. 4701–11.

53. Duran, J.M., et al., Unconventional secretion of Acb1 is mediated by autophagosomes. J Cell Biol, 2010. 188(4): p. 527–36.

54. Saerens, D., et al., Identification of a universal VHH framework to graft non-canonical antigen-binding loops of camel single-domain antibodies. J Mol Biol, 2005. 352(3): p. 597–607.

55. Aguilar, G., et al., Using Nanobodies to Study Protein Function in Developing Organisms. Antibodies (Basel), 2019. 8(1).

56. Rothbauer, U., et al., A versatile nanotrap for biochemical and functional studies with fluorescent fusion proteins. Mol Cell Proteomics, 2008. 7(2): p. 282–9.

57. Ressurreicao, M., S. Warrington, and D. Strutt, Rapid Disruption of Dishevelled Activity Uncovers an Intercellular Role in Maintenance of Prickle in Core Planar Polarity Protein Complexes. Cell Rep, 2018. 25(6): p. 1415–1424 e6.

58. Thoreen, C.C., et al., An ATP-competitive mammalian target of rapamycin inhibitor reveals rapamycin-resistant functions of mTORC1. J Biol Chem, 2009. 284(12): p. 8023–32.

59. Stolz, A., et al., Fluorescence-based ATG8 sensors monitor localization and function of LC3/GABARAP proteins. EMBO J, 2017. 36(4): p. 549–564.

60. Kirisako, T., et al., The reversible modification regulates the membrane-binding state of Apg8/Aut7 essential for autophagy and the cytoplasm to vacuole targeting pathway. J Cell Biol, 2000. 151(2): p. 263–76.

61. Kraft, L.J., et al., Nuclear LC3 Associates with Slowly Diffusing Complexes that Survey the Nucleolus. Traffic, 2016. 17(4): p. 369–99.

62. Klionsky, D.J., et al., Guidelines for the use and interpretation of assays for monitoring autophagy (4th edition)(1). Autophagy, 2021. 17(1): p. 1–382.

63. Jeong, J.H., D.K. Lee, and Y.H. Jo, Cholinergic neurons in the dorsomedial hypothalamus regulate food intake. Mol Metab, 2017. 6(3): p. 306–312.

64. Davis, L.I., The nuclear pore complex. Annu Rev Biochem, 1995. 64: p. 865–96.

65. Kaffman, A. and E.K. O’Shea, Regulation of nuclear localization: a key to a door. Annu Rev Cell Dev Biol, 1999. 15: p. 291–339.

66. la Cour, T., et al., analysis and prediction of leucine-rich nuclear export signals. Protein Eng Des Sel, 2004. 17(6): p. 527–36.

67. Kosugi, S., et al., Design of peptide inhibitors for the importin alpha/beta nuclear import pathway by activity-based profiling. Chem Biol, 2008. 15(9): p. 940–9.

68. Kosugi, S., et al., Systematic identification of cell cycle-dependent yeast nucleocytoplasmic shuttling proteins by prediction of composite motifs. Proc Natl Acad Sci U S A, 2009. 106(25): p. 10171–6.

69. Kosugi, S., et al., Six classes of nuclear localization signals specific to different binding grooves of importin alpha. J Biol Chem, 2009. 284(1): p. 478–85.

70. Jura, N., et al., Differential modification of Ras proteins by ubiquitination. Molecular Cell, 2006. 21(5): p. 679–687.

71. Xu, L., et al., Feedback regulation of Ras signaling by Rabex-5-mediated ubiquitination. Curr Biol, 2010. 20(15): p. 1372–7.

72. Nakagawa, T. and K. Nakayama, Protein monoubiquitylation: targets and diverse functions. Genes Cells, 2015. 20(7): p. 543–62.

73. Mahajan, R., et al., A small ubiquitin-related polypeptide involved in targeting RanGAP1 to nuclear pore complex protein RanBP2. Cell, 1997. 88(1): p. 97–107.

74. Speese, S.D., et al., Nuclear envelope budding enables large ribonucleoprotein particle export during synaptic Wnt signaling. Cell, 2012. 149(4): p. 832–46.

75. Wyant, G.A., et al., NUFIP1 is a ribosome receptor for starvation-induced ribophagy. Science, 2018. 360(6390): p. 751-758.

76. Geminard, C., E.J. Rulifson, and P. Leopold, Remote control of insulin secretion by fat cells in Drosophila. Cell Metab, 2009. 10(3): p. 199–207.

77. Gendrin, M., et al., Functional analysis of PGRP-LA in Drosophila immunity. PLoS One, 2013. 8(7): p. e69742.

78. Maitra, U., et al., Innate immune responses to paraquat exposure in a Drosophila model of Parkinson’s disease. Sci Rep, 2019. 9(1): p. 12714.

79. Masson, F., et al., Dual proteomics of Drosophila melanogaster hemolymph infected with the heritable endosymbiont Spiroplasma poulsonii. PLoS One, 2021. 16(4): p. e0250524.

80. Valanne, S., et al., Immune-inducible non-coding RNA molecule lincRNA-IBIN connects immunity and metabolism in Drosophila melanogaster. PLoS Pathog, 2019. 15(1): p. e1007504.

81. Shaw, P.J., et al., Stress response genes protect against lethal effects of sleep deprivation in Drosophila. Nature, 2002. 417(6886): p. 287-91.

82. Ekengren, S. and D. Hultmark, A family of Turandot-related genes in the humoral stress response of Drosophila. Biochem Biophys Res Commun, 2001. 284(4): p. 998–1003.

83. Agaisse, H., et al., Signaling role of hemocytes in Drosophila JAK/STAT-dependent response to septic injury. Dev Cell, 2003. 5(3): p. 441–50.

84. Ayres, J.S. and D.S. Schneider, The role of anorexia in resistance and tolerance to infections in Drosophila. PLoS Biol, 2009. 7(7): p. e1000150.

85. Barajas-Azpeleta, R., et al., Antimicrobial peptides modulate long-term memory. PLoS Genet, 2018. 14(10): p. e1007440.

86. Kim, K.A., et al., effect of dermcidin, an antimicrobial peptide, on body fat mobilization in normal mice. J Endocrinol, 2008. 198(1): p. 111–8.

87. Cardoso, F., et al., Neuro-mesenchymal units control ILC2 and obesity via a brain-adipose circuit. Nature, 2021. 597(7876): p. 410-414.

88. Vaziri, A., et al., Persistent epigenetic reprogramming of sweet taste by diet. Sci Adv, 2020. 6(46).

89. Mattila, J., et al., Mondo-Mlx Mediates Organismal Sugar Sensing through the Gli-Similar Transcription Factor Sugarbabe. Cell Rep, 2015. 13(2): p. 350–64.

90. Havula, E. and V. Hietakangas, Glucose sensing by ChREBP/MondoA-Mlx transcription factors. Semin Cell Dev Biol, 2012. 23(6): p. 640–7.

91. Kokki, K., et al., Metabolic gene regulation by Drosophila GATA transcription factor Grain. PLoS Genet, 2021. 17(10): p. e1009855.

92. Jacomin, A.C., et al., Regulation of Expression of Autophagy Genes by Atg8a-Interacting Partners Sequoia, YL-1, and Sir2 in Drosophila. Cell Rep, 2020. 31(8): p. 107695.

93. Knight, Z.A., et al., Hyperleptinemia is required for the development of leptin resistance. PloS one, 2010. 5(6): p. e11376.

94. Brooks, J.F., 2nd, et al., The microbiota coordinates diurnal rhythms in innate immunity with the circadian clock. Cell, 2021. 184(16): p. 4154–4167 e12.

95. Vaisse, C., et al., Leptin activation of Stat3 in the hypothalamus of wild-type and ob/ob mice but not db/db mice. Nat Genet, 1996. 14(1): p. 95–7.

96. Takats, S., et al., Investigating Non-selective Autophagy in Drosophila. Methods Mol Biol, 2019. 1880: p. 589–600.

97. Cai, X.T., et al., Gut cytokines modulate olfaction through metabolic reprogramming of glia. Nature, 2021. 596(7870): p. 97-102.

98. Chakrabarti, S., et al., Remote Control of Intestinal Stem Cell Activity by Haemocytes in Drosophila. PLoS Genet, 2016. 12(5): p. e1006089.

99. Brankatschk, M. and S. Eaton, Lipoprotein particles cross the blood-brain barrier in Drosophila. J Neurosci, 2010. 30(31): p. 10441–7.

100. Lee, P.T., et al., A gene-specific T2A-GAL4 library for Drosophila. Elife, 2018. 7.

101. Verboon, J.M., et al., Wash interacts with lamin and affects global nuclear organization. Curr Biol, 2015. 25(6): p. 804–810.

102. Kondo, S. and R. Ueda, Highly improved gene targeting by germline-specific Cas9 expression in Drosophila. Genetics, 2013. 195(3): p. 715–21.

103. Ro, J., Z.M. Harvanek, and S.D. Pletcher, FLIC: high-throughput, continuous analysis of feeding behaviors in Drosophila. PLoS One, 2014. 9(6): p. e101107.

104. Dobin, A., et al., STAR: ultrafast universal RNA-seq aligner. Bioinformatics, 2013. 29(1): p. 15–21.

105. Wang, L., S. Wang, and W. Li, RSeQC: quality control of RNA-seq experiments. Bioinformatics, 2012. 28(16): p. 2184–5.

106. Liao, Y., G.K. Smyth, and W. Shi, featureCounts: an efficient general purpose program for assigning sequence reads to genomic features. Bioinformatics, 2014. 30(7): p. 923–30.

107. Robinson, M.D. and A. Oshlack, A scaling normalization method for differential expression analysis of RNA-seq data. Genome Biol, 2010. 11(3): p. R25.

